# Distinct STRIPAK subunits drive conserved and subunit-specific signaling programs in *Cryptococcus neoformans*

**DOI:** 10.64898/2026.01.05.697761

**Authors:** Patricia P. Peterson, Sarah Croog, Yeseul Choi, Jin-Tae Choi, Yong-Sun Bahn, Joseph Heitman

**Affiliations:** Department of Molecular Genetics and Microbiology, Duke University Medical Center, Durham, North Carolina, USA; Department of Biotechnology, College of Life Science and Biotechnology, Yonsei University, Seoul, Republic of Korea

**Author notes:** Corresponding author (JH).

## Abstract

The striatin-interacting phosphatase and kinase (STRIPAK) complex is a conserved PP2A-associated signaling hub that integrates kinase-phosphatase networks, yet its roles in human fungal pathogens remain poorly defined. Here, we dissected STRIPAK functions in the opportunistic pathogen *Cryptococcus neoformans* by combining genetic, genomic, virulence, and phosphoproteomic analyses across mutants lacking individual STRIPAK subunits. Loss of the core STRIPAK components via *PPH22, FAR8, FAR9,* or *FAR11* mutations caused severe defects in growth, stress adaptation, cell-cycle progression, and morphogenesis, accompanied by widespread aneuploidy and genome instability. In murine infection models, *far11*Δ strains were avirulent, whereas *far9*Δ mutants caused delayed but ultimately fatal disease and underwent host-associated genome remodeling, with recovered isolates exhibiting chromosome 11 amplification despite no consistent *in vitro* fitness advantage. In contrast, deletion of *MOB3* produced a hypervirulent phenotype. *mob3*Δ cells exhibited enhanced transmigration across an i*n vitro* blood-brain barrier model, increased survival in macrophages, and generated small-cell morphotypes, features associated with increased dissemination. Phosphoproteomic profiling revealed extensive and overlapping phosphorylation changes among core STRIPAK mutants, affecting pathways involved in signaling, cytoskeletal and cell-cycle control, chromatin regulation, RNA metabolism, and stress responses. Conversely, *mob3*Δ mutants displayed a smaller, largely distinct phosphoproteomic signature. Network and functional enrichment analyses highlighted STRIPAK-dependent regulation of TORC2-associated signaling, MAPK/GTPase signaling, autophagy, nuclear transport, RNA processing, DNA replication, and ribosome biogenesis. Together, these findings establish STRIPAK as a coordinator of genome stability, morphological plasticity, stress adaptation, and virulence in *C. neoformans*, and demonstrate that individual STRIPAK subunits drive shared yet divergent signaling outputs that shape host-pathogen interactions.

**Importance:** Fungal pathogens must rapidly adapt their growth, morphology, and stress responses to survive within the host, requiring precise coordination of cellular signaling pathways. The conserved striatin-interacting phosphatase and kinase (STRIPAK) complex controls key developmental programs in eukaryotes, but its roles in fungal pathogenesis are not fully defined. We previously showed that STRIPAK is important for genome stability, development, and virulence in the opportunistic human fungal pathogen *Cryptococcus neoformans*. Here, we define how individual STRIPAK subunits differentially regulate fungal morphogenesis, genome plasticity, host adaptation, and virulence, revealing both shared and subunit-specific functions within this conserved signaling complex. Core STRIPAK mutants exhibit severe growth and stress-response defects and attenuation of virulence, whereas loss of the Mob3 subunit promotes hypervirulence by enhancing dissemination and persistence within the host. Phosphoproteomic profiling reveals that individual STRIPAK components exert shared yet distinct control over phosphorylation networks that shape host-pathogen interactions, establishing STRIPAK as a central signaling hub and a potential target for antifungal intervention.

## Introduction

Protein phosphorylation is a pervasive regulatory mechanism across eukaryotic organisms, governing fundamental processes such as growth, replication, and cellular differentiation in response to internal and external cues. The reversible addition of phosphate groups by protein kinases and their removal by phosphatases enables precise, dynamic control of protein activity and signaling flux. Together, these opposing enzymatic actions fine-tune signal transduction pathways that control cell cycle progression, metabolism, stress responses, transcription, and translation [1]. Dysregulated phosphorylation is implicated in numerous human diseases, including cancers and neurodegenerative disorders [2]. Accordingly, the coordinated balance between kinase and phosphatase activities is essential for maintaining cellular homeostasis.

The majority of eukaryotic protein phosphorylation occurs on serine and threonine residues, and among the phosphatases that reverse these modifications, protein phosphatase 2A (PP2A) is one of the most abundant and evolutionarily conserved [1]. Unlike kinases, whose substrate specificity is largely encoded within their catalytic domains, the functional diversity of PP2A arises from its ability to associate with a wide array of regulatory subunits. Assembly of distinct PP2A holoenzymes generates numerous isoforms with unique substrate specificities, subcellular localizations, and biological functions, allowing a single catalytic enzyme to govern diverse signaling pathways [3, 4]. Through these regulatory interactions, PP2A can directly modulate kinase cascades by dephosphorylating key kinases and signaling intermediates, positioning it as a central integrator of cellular signaling networks [5].

One prominent PP2A regulatory module is the striatin-interacting phosphatase and kinase (STRIPAK) complex, which incorporates the PP2A heterotrimeric holoenzyme, comprising the catalytic (C), scaffold (A), and a regulatory (B) subunit, also known as Striatin, together with additional adaptor proteins to form a distinct signaling scaffold [6-8]. Since its discovery, STRIPAK complexes have been identified across eukaryotes, from yeasts and filamentous fungi to *Drosophila* and mammals, where they play conserved roles in cytoskeletal organization, morphogenesis, stress responses, and developmental regulation [8-14]. Although the core components of STRIPAK are broadly conserved, many species encode multiple paralogs of individual subunits that retain shared functional domains, reflecting evolutionary diversification while preserving core signaling functions [15, 16]. Notably, the cellular processes regulated by STRIPAK are governed by highly interconnected signaling networks, highlighting STRIPAK’s central role as an organizing hub that coordinates kinase-phosphatase signaling during development and environmental adaptation.

In fungi, STRIPAK complexes have been most extensively studied in ascomycetous yeasts and filamentous species, including plant pathogens, where they play central roles in coordinating development, polarity, and environmental sensing. Genetic and biochemical studies across diverse fungal systems, including *Schizosaccharomyces pombe*, *Neurospora crassa, Sordaria macrospora*, and multiple filamentous plant pathogens, have demonstrated that STRIPAK regulates hyphal growth, septation, sexual reproduction, stress responses, and virulence-associated morphotypes [17]. In these organisms, STRIPAK functions at the intersection of MAPK signaling, cytoskeletal organization, and nuclear processes, acting as a scaffold that integrates kinase cascades with PP2A-mediated dephosphorylation. Disruption of STRIPAK components in plant-pathogenic fungi frequently results in severe developmental defects and impaired pathogenicity, underscoring the importance of this complex in fungal adaptation to host-associated environments [18-23]. Consistent with these roles, large-scale phosphoproteomic analyses in filamentous fungi have identified STRIPAK-dependent phosphorylation targets linked to vegetative growth and sexual development [24, 25]. Despite these advances in ascomycete fungi, far less is known about STRIPAK function in basidiomycetes or in human fungal pathogens.

*Cryptococcus neoformans* is a basidiomycete yeast and opportunistic human fungal pathogen that causes life-threatening meningoencephalitis, particularly in immunocompromised individuals, and is a leading cause of mortality among HIV/AIDS patients worldwide [26]. Establishing infection requires *C. neoformans* to dynamically balance cell proliferation with metabolic and morphological plasticity while adapting to diverse host environments, including the lung and central nervous system (CNS) [27]. Key virulence traits, such as thermotolerance, polysaccharide capsule production, titan cell formation, and the generation of small “seed” cells, are tightly linked to signaling pathways that regulate cell-cycle progression, stress responses, and cellular architecture [28, 29]. During infection, *C. neoformans* undergoes morphological transitions that promote survival and dissemination. Titan cells are enlarged polyploid cells that promote immune evasion, whereas seed cells are small morphotypes that facilitate dissemination to extrapulmonary tissues, including the central nervous system. Genome plasticity, including DNA mutation, aneuploidy, and polyploidization, has also emerged as an important adaptive strategy during infection, enabling rapid phenotypic diversification under host-imposed stresses [30-32]. Despite the central role of signaling networks in coordinating these processes, the contribution of STRIPAK-mediated PP2A signaling to *C. neoformans* biology is underexplored. Our previous work demonstrated that individual STRIPAK components regulate growth, genome stability, development, virulence-associated traits, and pathogenicity [33, 34]. However, how this conserved complex integrates signaling pathways at a systems level, and whether individual subunits exert shared or divergent functions, remains unexplored. Addressing this gap is essential for understanding how conserved phosphatase signaling networks are repurposed to control pathogenicity in human fungal pathogens.

In *C. neoformans*, the STRIPAK complex is predicted to comprise the PP2A scaffold Tpd3, the catalytic subunit Pph22, regulatory subunit and striatin homolog Far8, adaptor protein Far11, the tail-anchored membrane protein Far9, and the MOB-family subunit Mob3, forming a conserved multiprotein signaling assembly [15, 33]. In this study, we define the functions of previously uncharacterized STRIPAK components Far9 and Far11 and further investigate the basis of Mob3-dependent hypervirulence. We show that *FAR9* and *FAR11* are required for high-temperature growth, stress adaptation, and virulence-associated traits in *C. neoformans*. Despite severe growth defects, deletion of the *FAR9* or *FAR11* gene led to dramatic increases in cell size and widespread genome duplication, resulting in polyploidization. Whereas *far11*Δ mutants were avirulent, loss of *FAR9* caused delayed but ultimately fatal infection, accompanied by pronounced morphological heterogeneity within host tissues, altered cell wall organization, dissemination beyond the lungs and central nervous system, and evidence of genome evolution during infection.

In contrast, deletion of the gene encoding STRIPAK subunit *MOB3* resulted in a hypervirulent phenotype. Mechanistic analyses revealed that *mob3*Δ mutant cells exhibited enhanced transmigration across an in vitro blood–brain barrier model, increased survival within macrophages, and formed reduced cell body sizes under host-mimicking conditions and during pulmonary infection, features that may facilitate dissemination to the central nervous system. Finally, we employed a systems-level phosphoproteomic approach to characterize STRIPAK-dependent signaling across mutants lacking the PP2A catalytic subunit Pph22, regulatory subunit Far8/Striatin, or STRIPAK components Far11 and Mob3. These analyses revealed extensive remodeling of phosphorylation networks involved in signal transduction, cell-cycle regulation, cytoskeletal organization, and transcriptional and translational control. Protein interaction network analysis further identified STRIPAK-regulated modules spanning kinase signaling, RNA processing, ribosome biogenesis, autophagy, and DNA replication and maintenance. Notably, multiple STRIPAK mutants exhibited recurrent phosphorylation changes in proteins associated with TORC2 signaling and lipid homeostasis, including Efr3, Mss4, Pkh2, and Ypk1, as well as regulators of actin cytoskeleton organization and endocytic trafficking.

## Results

### Loss of the *FAR9* or *FAR11* gene causes broad defects in growth, stress resistance, and sexual development

Deletion of genes encoding core STRIPAK components and PP2A subunits Pph22 or Far8 causes broad fitness defects in *C. neoformans*, including sensitivity to temperature, nutrient, and cell wall stresses and impaired sexual development [33, 34]. In contrast, deletion of the STRIPAK component Mob3 results in increased thermotolerance, enhanced stress tolerance, and hypervirulence [33]. To further define the roles of the remaining conserved STRIPAK components, we focused on Far9, a tail-anchored membrane protein proposed to recruit STRIPAK to cellular membranes, and Far11, the central scaffolding subunit that coordinates assembly and organization of the STRIPAK complex [12, 35-37]. Although their primary sequences are highly divergent, multi-sequence alignments reveal conservation of key domains across fungi and humans, consistent with their assembly into a conserved STRIPAK complex (Figure S1). To determine whether these conserved STRIPAK subunits similarly contribute to fitness and development in *C. neoformans*, we deleted the genes encoding *FAR9* or *FAR11* in the H99α background (Figure S2) and assessed growth under diverse stress conditions (Figure 1A, S3A). Both *far9*Δ and *far11*Δ mutants displayed subtle growth defects on YPD at 30°C, but exhibited markedly reduced growth at 37°C and near-complete loss of growth at 25°C. The mutants showed almost no growth on nutrient-limiting RPMI medium and were hypersensitive to the calcineurin inhibitors FK506 and cyclosporine A, consistent with our previous findings that STRIPAK function becomes essential when calcineurin signaling is inhibited [33]. *far9*Δ and *far11*Δ strains were also highly sensitive to ionic and cell wall stress, including KCl, NaCl, SDS, calcofluor white, caffeine, and Congo red. Growth on medium supplemented with 1 M sorbitol resulted in a modest improvement in colony size but did not restore wild-type growth, suggesting that impaired cell integrity contributes to, but does not fully explain, the observed growth defects.

**Figure 1.**
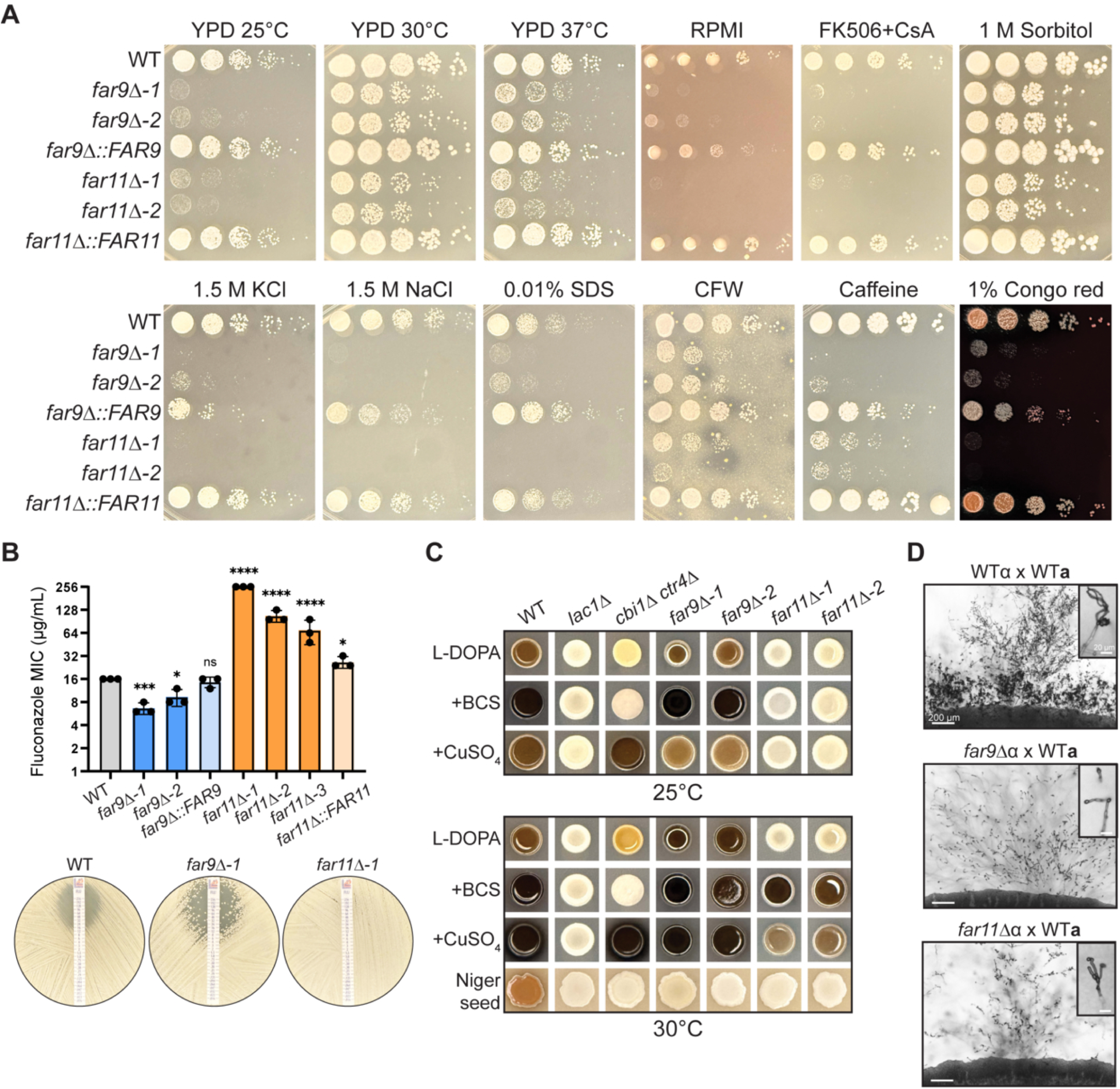
STRIPAK complex *far9*Δ and *far11*Δ mutants exhibit significant defects in growth, stress response, and sexual development. (A) WT (H99α) and isogenic *far9*Δ and *far11*Δ strains, and their respective genetically complemented strains, were serially diluted and plated on YPD at 25°C, 30°C, and 37°C; RPMI; and YPD supplemented with 1 μg/mL FK506/100 μg/mL cyclosporine A (CsA), 1 M sorbitol, 1.5 M potassium chloride (KCl), 1.5 M sodium chloride (NaCl), 0.01% SDS, 1.5 mg/mL calcofluor white (CFW), 0.5 mg/mL caffeine, or 1% Congo red—each at 30°C. Images were taken after 2 days of incubation. (B) Fluconazole Etest to analyze drug susceptibility in WT (H99α), *far9*Δ, and *far11*Δ strains. Cells were grown in an overnight culture in YPD to saturation, then spread onto YPD plates before adding FLC Etest strips. Plates were incubated at 30°C and images were taken after 48 hours. Quantification of MIC values from three independent biological replicates is shown. (C) WT (H99α), *far9*Δ, and *far11*Δ were serially diluted and plated onto Niger seed or L-DOPA agar medium to induce melanin production. For L-DOPA assays, medium was supplemented with either bathocuproine disulfonate (BCS) or CuSO_4_ where indicated to assess the effects of copper availability. Plates were incubated at 25°C or 30°C as indicated and imaged after 5 days. The *lac1*Δ mutant served as a melanin-deficient control, and the *cbi1*Δ *ctr4*Δ strain was included as a control for copper-dependent melanization. (D) Reduced mating efficiency of *far9*Δ and *far11*Δ. Cells were co-cultured with wild-type cells of the opposite mating type (KN99**a**) on MS plates. An H99α x KN99**a** cross served as a control. Images of mating patches and basidia were taken after 5 weeks’ incubation at room temperature.

To verify that the observed phenotypes were attributable to loss of *FAR9* or *FAR11*, complemented strains were generated and expression was confirmed by RT-qPCR (Fig. S2E, S2F, S3B, S3C). *far9*Δ*::FAR9* complemented strains expressed *FAR9* at levels comparable to or exceeding those observed in the wild type, whereas the *FAR11* complemented strain exhibited approximately two-fold higher expression that was not significantly different from wild type. Importantly, growth defects across these tested conditions were partially or fully restored in the complemented *far9*Δ*::FAR9 and far11*Δ*::FAR11* strains, confirming that the phenotypes result from loss of the respective STRIPAK components.

Given the pronounced cell wall and stress sensitivities of the *far9*Δ and *far11*Δ mutants, we next examined their responses to the ergosterol-targeting antifungal fluconazole (Figure 1B, S3D). Based on Etest assays, *far9*Δ mutants exhibited a lower MIC (8 to 12 μg/mL) relative to wild-type (16 μg/mL), indicating increased fluconazole susceptibility. However, despite the reduced MIC, *far9*Δ mutants produced many colonies within the zone of inhibition, consistent with enhanced tolerance. In contrast, *far11*Δ mutants showed near-complete growth along the fluconazole gradient (MIC 64 to 256 μg/mL), indicating fluconazole resistance. These phenotypes were partially or fully restored in the complemented strains. These findings suggest that Far9 and Far11 play distinct roles in the cellular responses to membrane stress and fluconazole exposure.

We next examined melanin production in *far9*Δ and *far11*Δ mutants, as melanin is a key virulence factor protecting *Cryptococcus* from environmental and host-derived stresses (Figure 1C, S3F, S3G). On Niger seed medium, only wild-type cells produced melanin, whereas *far9*Δ, *far11*Δ, and the melanin-defective control *lac1*Δ mutant remained non-pigmented. To assess whether copper availability influenced melanization, we tested pigmentation on L-DOPA medium supplemented with either copper sulfate or the copper chelator bathocuproine disulfonate (BCS) at both 25°C and 30°C. The *cbi1*Δ *ctr4*Δ strain, which is defective in high-affinity copper uptake and copper binding, was included as a control for copper-dependent melanization, producing pigment only in the presence of added copper and remaining non-pigmented with BCS [33, 38, 39]. Wild-type melanized robustly on all L-DOPA conditions tested, independent of copper availability or temperature. *far9*Δ mutants were likewise capable of melanization under all conditions tested. At 25°C, both wild type and *far9*Δ strains exhibited darker pigmentation on BCS-containing medium than on copper-supplemented medium. In contrast, *far11*Δ mutants failed to melanize under any condition at 25°C. At 30°C, however, they produced no pigment on L-DOPA alone but melanization was restored by both copper supplementation and chelation, with BCS producing the strongest pigmentation. The strain-specific melanization defects observed in *far9*Δ and *far11*Δ mutants were restored in the corresponding complemented strains (Figure S3E, S3F). Together, these findings suggest that STRIPAK influences melanization in a manner that is responsive to environmental copper conditions, although the underlying relationship appears more complex than a simple defect in copper acquisition.

STRIPAK is broadly conserved as a regulator of fungal sexual development [4,5]. To define the specific contributions of Far9 and Far11 to *C. neoformans* mating, we crossed the corresponding deletion mutants with a wild-type strain of the opposite mating type (KN99**a**) on MS medium and monitored mating structures over time (Figure 1D). The H99α × KN99**a** cross served as the wild-type control. Because *far9*Δ and *far11*Δ mutants exhibit impaired growth at room temperature, we employed two strategies to facilitate mating: (1) mixing mutant and wild-type cells at a 10:1 ratio, and (2) pre-growing the mutant alone on MS for three days before overlaying with wild-type cells. After six weeks, *far9*Δ x WT crosses produced extensive hyphal growth with large peripheral branches resembling the wild-type cross, whereas *far11*Δ x WT crosses generated markedly reduced hyphal development. Despite these differences, both mutants failed to complete sexual development: the basidia that formed were predominantly bald, with little to no spore production. Defects in sexual development were consistently observed across independent mutant clones and were restored by genetic complementation (Figure S3G). These findings demonstrate that Far9 and Far11 are required not only for robust hyphal formation but also for successful basidia maturation and sporulation.

### *far9*Δ and *far11*Δ mutations cause significant genome instability and cell cycle defects

Our previous work showed that STRIPAK complex mutants, including *pph22*Δ*, far8*Δ, and *mob3*Δ, frequently accumulate segmental and whole-chromosome aneuploidy in both haploid and heterozygous diploid backgrounds, accompanied by defects in cell-cycle progression [33]. To determine whether *FAR9* and *FAR11* contribute similarly to genome maintenance, we performed whole-genome sequencing of *far9*Δ and *far11*Δ mutant strains and analyzed read-depth coverage across all chromosomes (Figure 2A, S4A, S4B). Sequencing confirmed the expected gene deletions and did not identify additional high-confidence sequence variants among isolates. Normalized read-depth profiles showed increases in coverage approaching two-fold across multiple chromosomes relative to the euploid H99α control, consistent with whole-chromosome aneuploidy. The extent of genome imbalance varied between the two mutants: *far9*Δ strains exhibited only mild aneuploidy, predominantly affecting chromosomes 2 and 11; whereas *far11*Δ mutants displayed widespread aneuploidy, with increased copy number affecting approximately half of the14 chromosomes. The specific chromosomes exhibiting copy-number variation differed among the independently generated isolates. Importantly, restoration of *FAR11* returned chromosome copy number profiles to a euploid state whereas the *FAR9* complemented strain retained elevated chromosome 11 copy number, although the extent of aneuploidy was reduced relative to the parental *far9*Δ-1 strain (Figure 2A, S4B). These findings indicate that *far9*Δ and *far11*Δ mutants exhibit chromosome copy number variation and genome instability, with *far11*Δ mutants displaying extensive chromosomal aneuploidy consistent with progressive genome duplication toward a diploid state.

**Figure 2.**
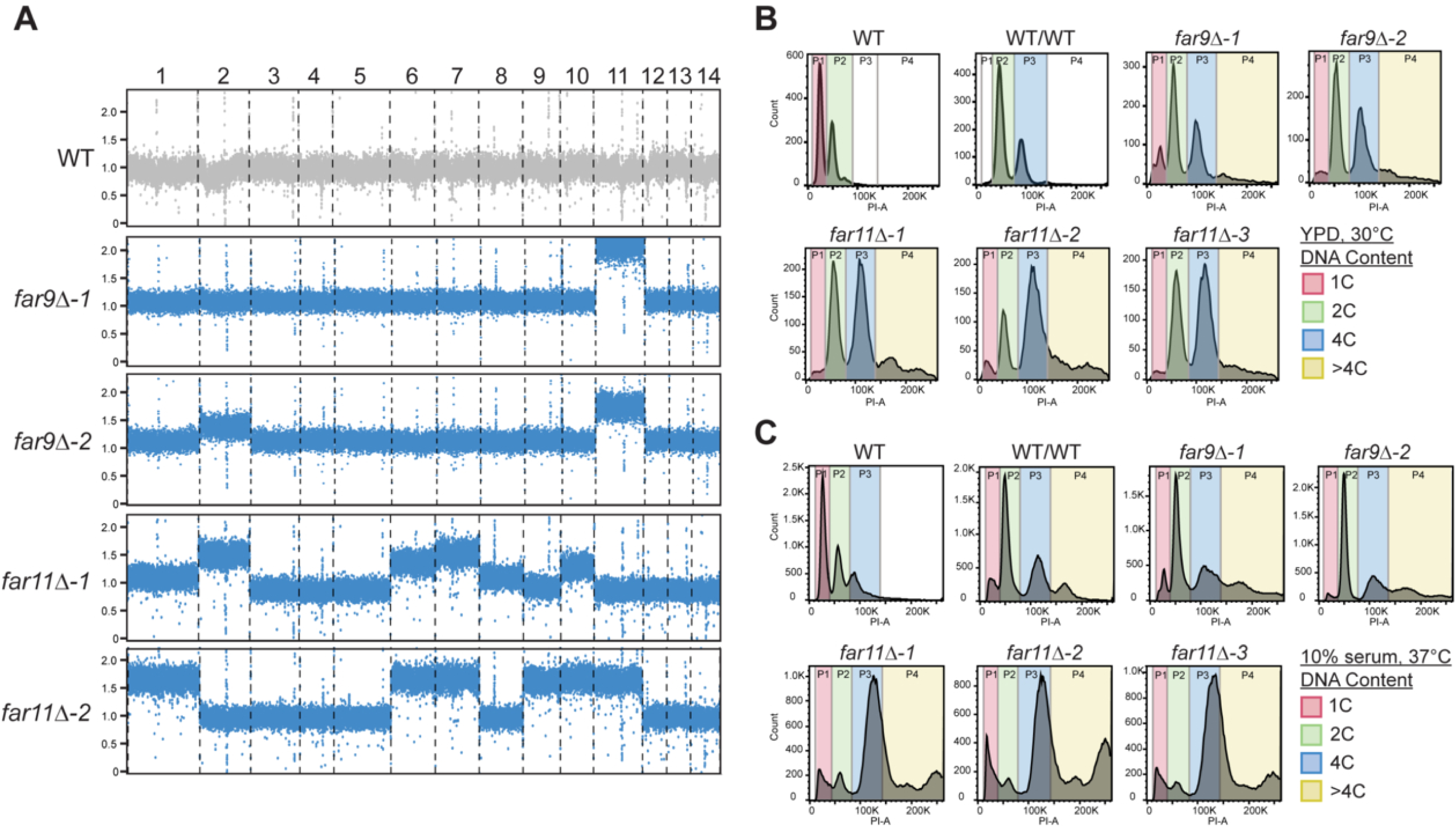
Loss of Far9 or Far11 causes genome instability. (A) Read depth analyses from whole-genome sequencing of the H99α (WT) parental strain and *far9*Δ and *far11*Δ strains. Read coverage was normalized to the genome-wide median, such that euploid chromosomes exhibit approximately 1X coverage. Both mutants display chromosome copy-number variation, with *far11*Δ isolates showing more extensive aneuploidy than *far9*Δ isolates. (B) Flow-cytometric analysis of DNA content in WT, diploid WT/WT, *far9*Δ, and *far11*Δ strains grown in YPD at 30°C. Cells were stained with propidium iodide and analyzed by flow cytometry. Colored gates indicate populations with 1C, 2C, 4C, and >4C DNA content. H99α (haploid) and WT/WT (KN99**a**/KN99α, diploid) served as ploidy controls. (C) When grown in 10% fetal bovine serum (FBS) at 37°C, both *far9*Δ and *far11*Δ strains exhibited altered DNA-content distributions relative to WT controls, including expanded 4C and >4C populations.

We next applied fluorescence-activated cell sorting (FACS) to assess DNA content in *far9*Δ and *far11*Δ mutants under both standard and host-mimicking culture conditions. When grown in YPD at 30°C, both mutants displayed altered DNA-content distributions characterized by expanded 4C and >4C populations relative to the haploid wild type (Figure 2B). Notably, both *far9*Δ and *far11*Δ strains exhibited an expanded fraction of cells with ≥4C DNA content, indicating the emergence of a polyploid subpopulation. In YPD, the combined P3/P4 (≥4C) populations represented 45-72% of mutant cells, compared with only 4% of the haploid control and 32% of the diploid control (Figure S4D).

To evaluate DNA content distributions under physiologically relevant conditions, we grew the strains in 10% fetal bovine serum (FBS) at 37°C, an established *in vitro* cue for titan cell induction [40], and repeated the FACS analysis (Figure 2C). Under serum conditions, the diploid control shifted to a higher proportion of ≥4C cells, and therefore more closely resembled the *far9*Δ profile, which remained relatively stable between YPD and serum. In contrast, *far11*Δ mutants exhibited a dramatic redistribution toward polyploid DNA content. The major *far11*Δ peak shifted beyond the 4C gate, with a substantial fraction of cells accumulating in the >4C population. In serum, the P4 population (>4C) represented approximately 24 to 43**%** of *far9*Δ and *far11*Δ cells, compared with only 4% and 16% in the haploid and diploid controls, respectively (Figure S4D).

To determine whether restoration of STRIPAK function could rescue these defects, we analyzed DNA content in the complemented strains under both YPD and serum conditions (Figure S4E). Although DNA-content profiles in YPD remained broadly similar to those observed in the parental mutants (Figure 2B, 2C), restoration of *FAR9* and *FAR11* substantially reduced the accumulation of cells with elevated DNA content under serum conditions. In particular, the expanded >4C populations observed in *far11*Δ mutants were largely eliminated following restoration of *FAR11*. These findings confirm that the aberrant DNA-content distributions are attributable to disruption of the respective STRIPAK components. Collectively, the genome sequencing and FACS analyses demonstrate that disruption of STRIPAK function leads to chromosome copy-number variation, altered DNA-content distributions, and defects in cell-cycle progression. Both *far9*Δ and *far11*Δ mutants formed heterogeneous populations containing cells with elevated genomic content, although the extent and distribution of these populations differed between strains and growth conditions.

### *far9*Δ and *far11*Δ mutant strains exhibit increased cell size under both nutrient-rich and nutrient-limiting conditions

Because changes in ploidy and cell-cycle progression often alter fungal morphology, we next assessed whether loss of *FAR9* or *FAR11* affects cell size or cellular architecture. We examined cell morphology in *far9*Δ and *far11*Δ mutants under nutrient-rich (YPD) and nutrient-limiting (RPMI) conditions at 30°C (Figure 3). In YPD, both mutants appeared substantially larger than wild-type cells and frequently contained prominent vacuoles occupying a large portion of the cell volume, features not observed in the wild type (Figure 3A). In RPMI, wild-type cells were slightly smaller than in YPD, whereas *far9*Δ and *far11*Δ mutant cells remained significantly larger with pronounced vacuoles (Figure 3B). India ink staining showed that capsule thickness was not increased in either mutant despite the larger cell size. Quantification revealed markedly broader cell-size distributions in both mutants (Figure 3C). In YPD, wild-type cells ranged from 3.4 to 7.9 μm, compared with 4.5 to 10.1 μm in *far9*Δ and 4.2 to 12.2 μm in *far11*Δ. In RPMI, wild-type cells ranged from 3.1 to 7.1 μm, compared with 3.2 to 15.4 μm in *far9*Δ and 3.3 to 13.5 μm in *far11*Δ. These broadened distributions reveal substantial morphological heterogeneity in both mutants, which was consistent across biological triplicates. The enlarged cell size of *far9*Δ and *far11*Δ mutants in YPD was accompanied by increased cell aggregation and rapid settling in liquid culture (Figure S5), suggestive of flocculation-like behavior. The presence of highly heterogeneous cell populations with a substantial fraction of cells >10 μm in diameter, together with the increased ≥4C DNA-content identified in our FACS analyses, suggests that *far9*Δ and *far11*Δ strains may exhibit features associated with titan cell development under both inducing and non-inducing conditions. These morphological and DNA content phenotypes are consistent with titan-associated traits previously described in *C. neoformans* [40].

**Figure 3.**
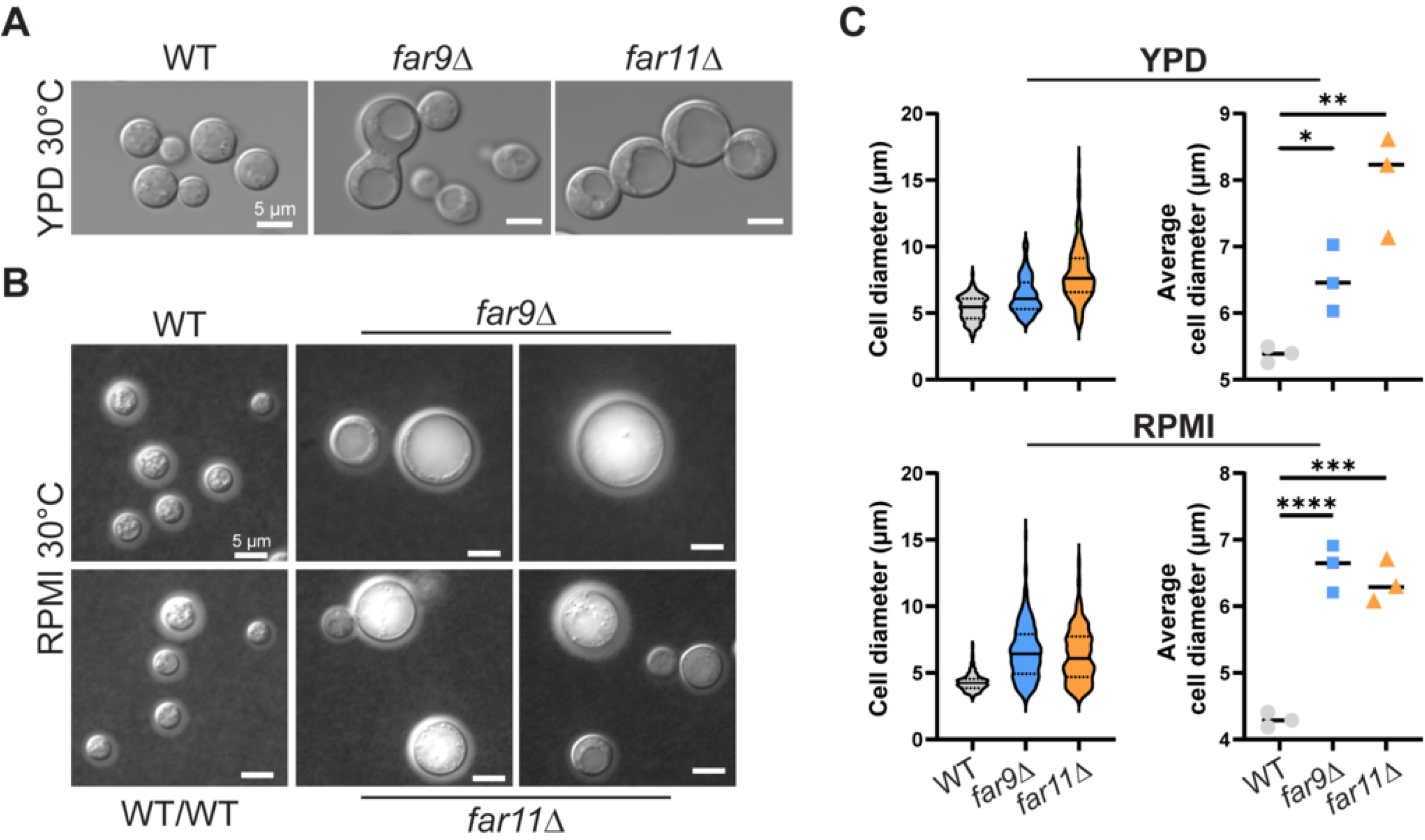
Far9 and Far11 promotes cell enlargement and morphological heterogeneity. (A) Differential interference contrast (DIC) microscopy images of WT, *far9*Δ, and *far11*Δ cells grown to mid-logarithmic phase in YPD liquid media at 30°C. Mutant strains exhibited markedly enlarged cell bodies and prominent intracellular vacuoles occupying a large proportion of the cell volume. (B) DIC and India ink staining images of WT, *far9*Δ, and *far11*Δ cells grown under capsule-inducing conditions in RPMI medium at 30°C. (C) Quantification of cell diameters across three biological replicates revealed significantly larger and more heterogeneous cell populations in the mutant strains. Images were analyzed with ImageJ/Fiji. Data are presented as scatter plots of individual cells (left) and biological replicate means (right).. Statistical significance was calculated using one-way ANOVA with Dunnett’s multiple comparisons test.

### STRIPAK mutants show reduced virulence and undergo genomic and morphological changes *in vivo*

The severe growth impairment of *far9*Δ and *far11*Δ mutants at 37°C and during nutrient stress suggested that these strains might be attenuated for virulence. Indeed, deletion of genes encoding core STRIPAK components Pph22 or Far8, which display similar stress-sensitive phenotypes, are fully avirulent in mice [33]. To assess the *in vivo* consequences of *FAR9* or *FAR11* loss, we infected A/J mice intranasally with wild-type, *far9*Δ, or *far11*Δ mutant strains and monitored for survival. As expected, *far11*Δ was completely avirulent. In contrast, and unexpectedly, mice infected with *far9*Δ exhibited delayed but ultimately severe disease progression. After a prolonged asymptomatic period, *far9*Δ-infected animals began to succumb rapidly beginning around day 40 post infection, resulting in near-complete mortality by day 80 when the experiment was terminated (Figure 4A). An independent infection experiment that included complemented *far9*Δ::*FAR9* and *far11*Δ::*FAR11* strains reproduced the virulence phenotypes observed for the mutants (Figure S6A). Restoration of *FAR11* fully rescued virulence, whereas the *far9*Δ::*FAR9* strain exhibited an intermediate phenotype characterized by delayed disease progression relative to wild type, but increased virulence compared with the *far9*Δ mutant.

**Figure 4.**
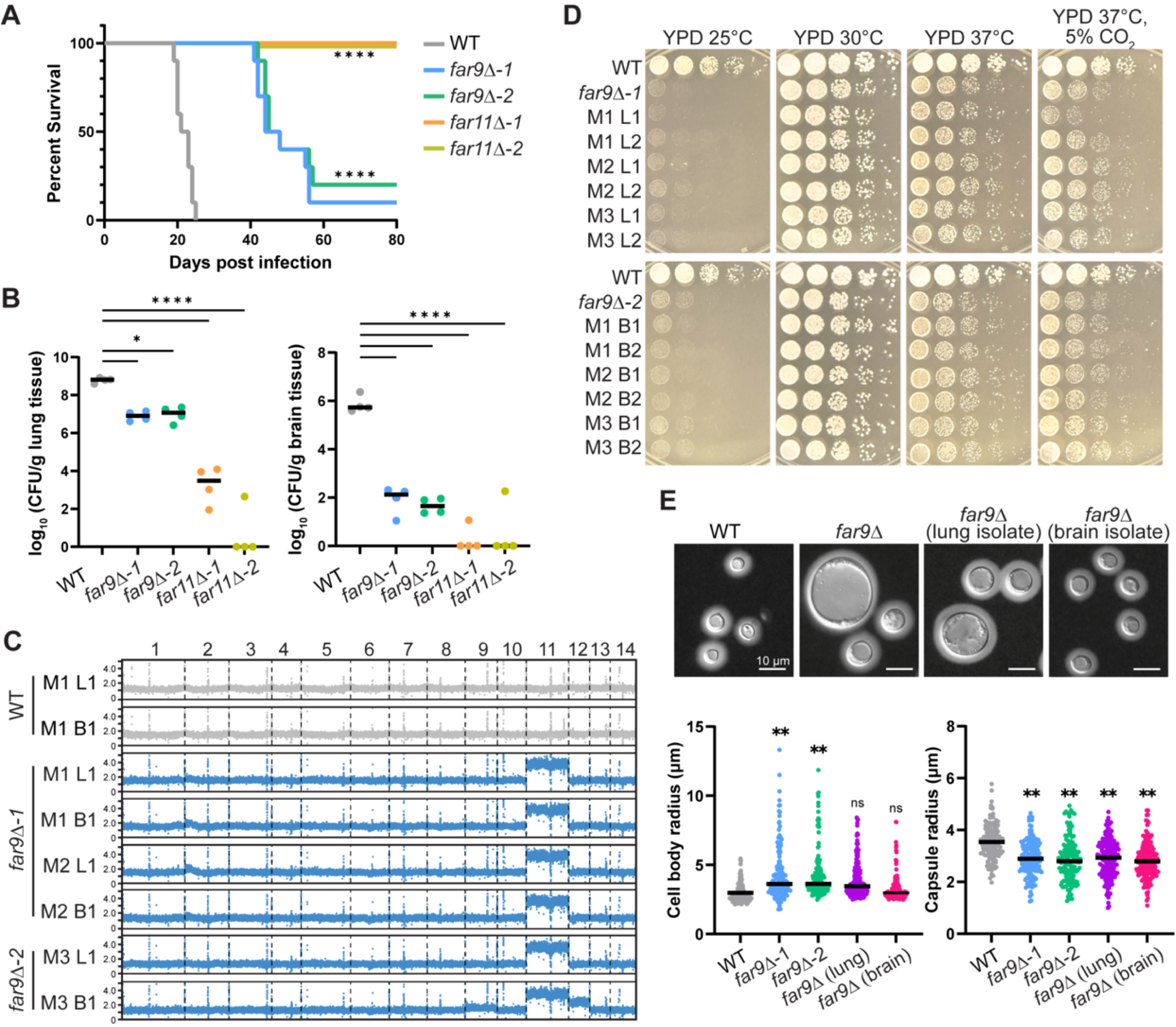
Virulence attenuation of *far9*Δ and *far11*Δ mutants and characterization of mouse-passaged *far9*Δ isolates. Equal numbers of male and female A/J mice were infected intranasally with 1 x 10^5^ cells of the indicated WT and isogenic mutant strains and analyzed for survival rate (n=10) and fungal burden (n=4). (A) Survival analysis of WT (H99α), *far9*Δ, and *far11*Δ infected mice. Mice were 5 weeks old at the time of infection. (B) CFUs per gram of lung and brain tissue recovered from organs harvested at 14 days post-infection (DPI). Statistical significance was assessed using one-way ANOVA with Dunnett’s multiple-comparisons testing. (C) Read-depth analysis from whole-genome sequencing of *far9*Δ isolates recovered from the indicated tissues of independent animals at terminal disease. All recovered isolates retained increased chromosome 11 copy number, whereas the chromosome 2 copy-number increase observed in the parental *far9*Δ strains (Figure 2A) was no longer detected. Isolate labels indicate mouse number (M) and tissue source (B, brain; L, lung). (D) Serial dilution spotting assays comparing growth of parental *far9*Δ strains and mouse-passaged isolates on YPD at 25°C, 30°C, 37°C, and 37°C with 5% CO_2_. Images were captured after 3 days of incubation. (E) Cell body and capsule measurements of WT, *far9*Δ, and *far9*Δ isolates recovered from mice. Cells were grown in 10% FBS at 37°C with 5% CO_2_ and stained with India ink. Measurements represent three biological replicates for WT and the parental *far9*Δ strain and independent recovered isolates from infected animals. Statistical significance was determined using biological replicate means rather than individual cell measurements and was assessed by one-way ANOVA with Tukey’s multiple-comparisons test.

To determine whether delayed mortality reflected impaired dissemination or altered disease kinetics, fungal burden was assessed in four animals per group at day 14 post infection. At this early time point, *far9*Δ mutant cells persisted in the lungs at significantly reduced levels compared with wild type yet were consistently recovered from the brains of all infected animals, albeit at low levels (Figure 4B). In contrast, *far11*Δ mutants exhibited greater attenuation, with markedly reduced lung colonization and brain dissemination detected in only 2 of 8 infected animals. These findings indicate that Far11 is required for establishment of infection, whereas loss of Far9 does not prevent early dissemination but results in a prolonged latent phase prior to eventual lethal disease.

To further investigate the unusual disease course of *far9*Δ infections, we performed an additional infection experiment and assessed fungal burden at both 7 and 14 days post infection (Figure S6B, S6C). Consistent with the initial survival studies, *far9*Δ mutants exhibited delayed disease progression, whereas the complemented strain displayed an intermediate virulence phenotype. At both time points, *far9*Δ fungal burdens remained significantly reduced relative to wild-type in the lungs and brain. Although complementation partially restored tissue colonization, fungal burdens remained lower than those observed for the wild-type. Together, these findings confirm that loss of *FAR9* impairs fungal proliferation *in vivo* while permitting persistence and eventual progression to lethal disease.

Because *far9*Δ mutants were initially predicted to be avirulent, we next sought to verify that the animals succumbing after day 40 were indeed infected with the original mutant strain. Single colonies were isolated at humane endpoints from the lungs and brains of three independently infected animals (derived from two independent *far9*Δ parental strains) and subjected to whole-genome sequencing. All isolates retained the *far9*Δ deletion, confirming that the fatal infections were caused by the mutant strain. Read-depth analyses revealed that chromosome 11 aneuploidy was maintained in all mouse-passaged isolates, whereas wild-type isolates recovered from infected animals remained euploid (Figure 4C). Flow cytometry analysis demonstrated that the recovered *far9*Δ isolates retained cell populations with elevated DNA-content relative to wild-type (Figure S7A). Unlike chromosome 11, the chromosome 2 copy-number increase observed in the parental *far9*Δ-2 strain (Figure 2A) was not maintained in the recovered isolates. One brain isolate additionally exhibited elevated chromosome 12 copy number. Together, these findings indicate that chromosome 11 aneuploidy is a stable feature of *far9*Δ populations during infection.

To determine whether *in vivo* passage altered the fitness of *far9*Δ mutants, we compared mouse-passaged isolates to the parental *far9*Δ strains under multiple growth conditions. Serial dilution spotting assays at 25⁰C, 30⁰C, 37⁰C, and 37⁰C in 5% CO_2_ revealed no obvious growth advantage in any of the recovered isolates (Figure 4D). The passaged strains remained highly sensitive to low temperature and failed to grow at 25⁰C, similar to the parental *far9*Δ mutants. Likewise, no improvements were observed on nutrient-limiting RPMI, on YPD supplemented with FK506 and cyclosporine A, or on melanin pigment-inducing medium (Figure S7B, S7C). Thus, despite prolonged persistence and disease progression *in vivo*, the recovered isolates did not exhibit obvious improvements in the *in vitro* phenotypes examined.

To further assess whether *in vivo* passage produced morphological or capsule-related adaptations, we examined India ink–stained cells grown in 10% serum at 37⁰C with 5% CO_2_, comparing wild type, the *far9*Δ parental strain, and isolates recovered from the lungs and brain (Figure 4E, S7D, S7E). Quantification revealed that the parental *far9*Δ strains had a significantly larger cell body radius and reduced capsule radius relative to wild type, consistent with the morphology phenotypes shown in Figure 3. Both lung- and brain-derived *far9*Δ isolates exhibited significantly smaller average cell body sizes than the parent strains, although enlarged cells remained present within the recovered populations. In contrast, capsule size remained significantly reduced relative to wild type and did not differ significantly between the parental and mouse-passaged *far9*Δ populations. These findings indicate that *in vivo* passage is associated with remodeling of *far9*Δ cell morphology, particularly cell body size, while the reduced capsule phenotype is maintained.

To characterize the morphology of *far9*Δ cells during infection, lungs collected at day 14 post-infection were homogenized and stained with calcofluor white (CFW) to visualize chitin in the fungal cell wall and distinguish fungal cells from host material (Figure 5A-C). In wild-type infections, *C. neoformans* cells displayed the expected yeast morphology, with occasional titan cells and pseudohyphal forms noted. WT cells consistently showed a smooth, continuous CFW-stained cell wall encircling each cell (Figure 5A). In contrast, *far9*Δ cells exhibited striking morphological abnormalities (Figure 5B, 5C). Mutant cells frequently formed irregular buds, enlarged cells, and elongated chains of connected cells, consistent with defects in cytokinesis and cell separation. CFW staining revealed discontinuous and uneven chitin deposition, with intense chitin accumulation at presumed division sites where daughter cells failed to separate. These features indicate severe defects in cell division and cell-wall organization during infection. *far9*Δ infections also produced giant cells exceeding 50 µm in diameter (Figure 5C). Notably, these enlarged cells exhibited fragmented CFW staining and aberrant chitin deposition (Figure 5B and 5C), rather than the continuous peripheral staining seen in wild type, suggesting substantial alterations in chitin organization and cell wall architecture during cell enlargement.

**Figure 5.**
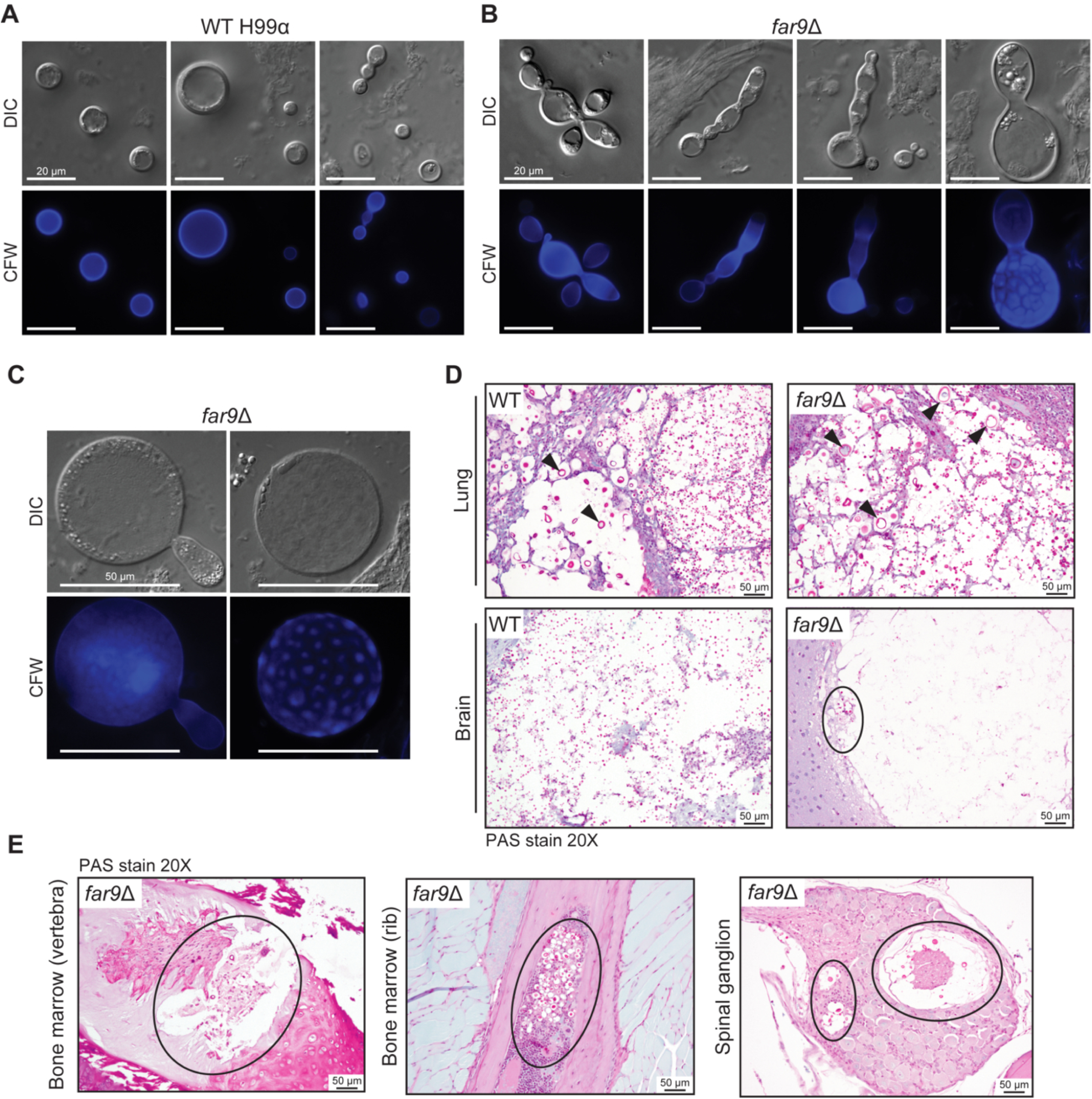
*far9*Δ mutant cells exhibit extensive morphological heterogeneity during infection. (A-C) Lung homogenates recovered at 14 days post infection (from experiment shown in Figure 4) were stained with calcofluor white (CFW) to observe chitin in the fungal cell wall. (A) WT type cells displayed predominantly yeast morphology with occasional enlarged cells and pseudohyphae and uniform chitin staining. (B) *far9*Δ cells show abnormal patterns of multi-budding, elongation, pseudohyphae, and enlarged cells, and uneven chitin deposition. (C) *far9*Δ can form giant titan cells with fragmented staining patterns of the cell wall. (D) Histopathological analysis of organs recovered when humane experimental endpoints were reached for WT- (day 21) and *far9*Δ- (day 48) infected animals. Lung and brain sections were stained with Periodic Acid–Schiff (PAS). Arrows indicate titan-like cells in lung tissue, which are larger and more pronounced in *far9*Δ infections. The circled region in the brain highlights the sparse *far9*Δ fungal cells within a representative brain lesion. Four animals were analyzed per group. (E) Representative PAS-stained sections showing *far9*Δ cells within vertebral and rib-associated bone marrow and spinal ganglia. Fungal cells were observed in these tissues only in *far9*Δ-infected animals and were not detected in WT infections.

To assess tissue pathology and fungal dissemination at terminal disease, lungs, brains, and additional tissues were collected at the time of sacrifice (day 21 for WT infections and day 48 for *far9*Δ). In the lungs, wild type infections displayed predominantly yeast morphologies with occasional titan-like cells, whereas *far9*Δ mutant infections exhibited pronounced morphological heterogeneity, characterized by giant cells, pseudohyphal and elongated forms, and smaller yeast-like cells (Figure 5D, S6D). In the brain, both WT- and *far9*Δ-infected animals exhibited cryptococcomas and associated tissue damage, indicating successful dissemination to the CNS. However, *far9*Δ fungal cells appeared less abundant and less extensively distributed within brain lesions than those observed in wild type infections (Figure 5D). Strikingly, fungal cells were detected within vertebral and rib-associated bone marrow as well as spinal ganglia in *far9*Δ-infected animals, sites not observed in wild type infections (Figure 5E). Together, these findings indicate that although *far9*Δ mutants exhibit reduced fungal burden early during infection, they are capable of persisting within the host, disseminating beyond the lungs and brain, and undergoing extensive morphological diversification during disease progression.

### *MOB3* deletion promotes hypervirulence through enhanced BBB crossing and altered cell morphology

Our previous studies showed that deletion of the STRIPAK subunit Mob3 results in a hypervirulent phenotype characterized by increased thermotolerance, enhanced melanin and capsule production, greater tissue infiltration during infection, and reduced host survival [33]. To further investigate the mechanistic basis of *mob3*Δ mutant hypervirulence, we employed an *in vitro* blood-brain barrier (BBB) model to quantify fungal transmigration across a monolayer of endothelial cells grown on a transwell membrane (Figure 6A). This assay has been widely applied in pathogenic *Cryptococcus* species to identify determinants of CNS invasion [41-44]. Independent *mob3*Δ isolates, generated from independent meiotic progeny described previously [33], exhibited significantly greater transmigration across the BBB compared with wild type, while trans-endothelial electrical resistance (TEER) remained unchanged following incubation (Figure 6B), indicating that increased transmigration was not due to disruption of endothelial barrier integrity.

**Figure 6.**
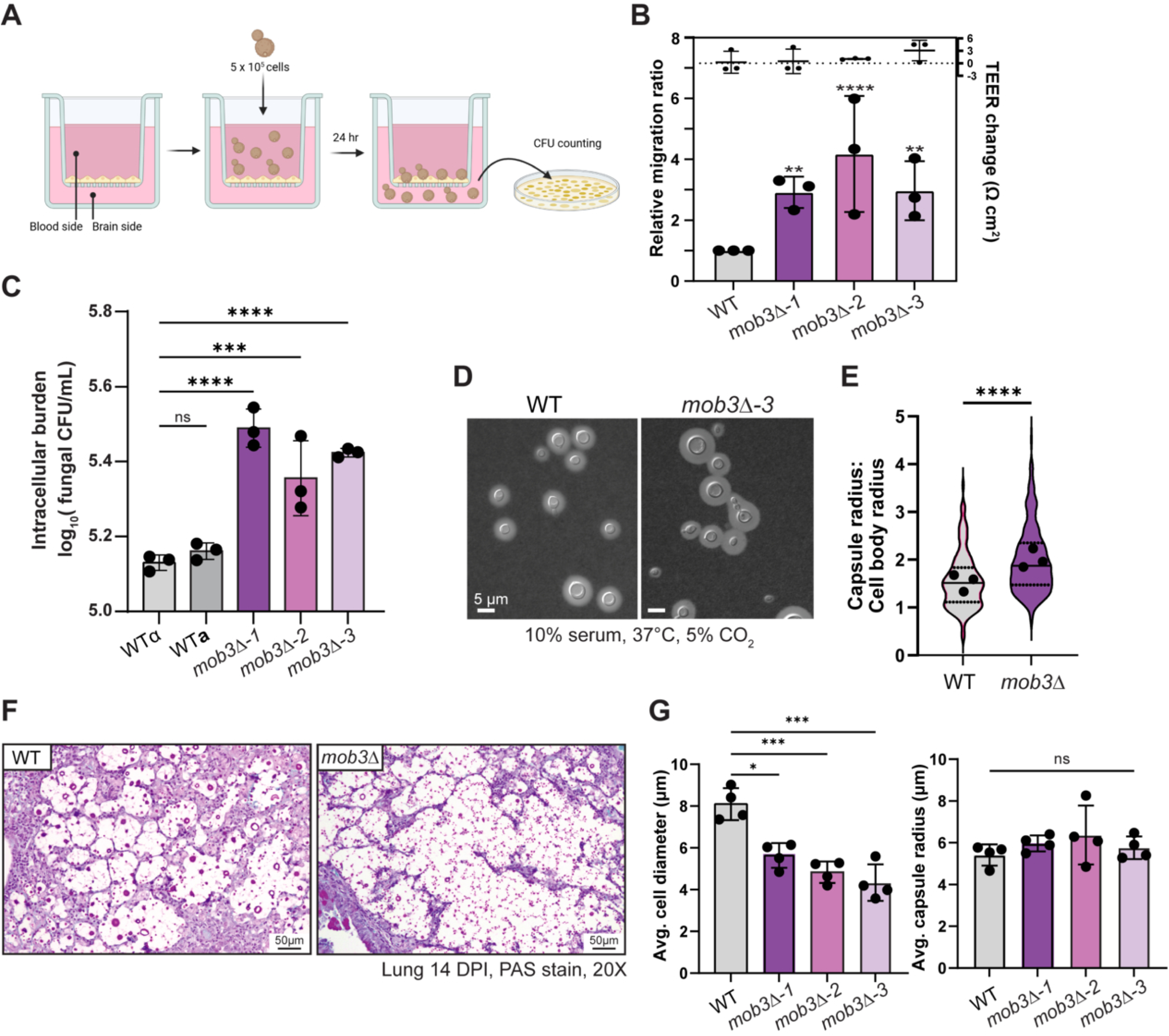
Hypervirulent *mob3*Δ strains exhibit enhanced blood-brain barrier (BBB) crossing, macrophage survival, and dissemination-associated morphologies. (A) Schematic diagram of the BBB transwell model and plating assay to determine rate of transmigration through the endothelium. (B) *mob3*Δ strains show significantly increased transmigration through the BBB compared to the wild-type. The TEER values remained relatively unchanged from the beginning to the end of the experiment, indicating that increased transmigration was not associated with disruption of endothelial barrier integrity. (C) Survival of WT and *mob3*Δ strains following 24 h co-incubation with THP-1 macrophages. Significantly more viable *mob3*Δ mutant cells were recovered, indicating enhanced intracellular survival and/or proliferation. (D) WT and *mob3*Δ cells were grown in 10% fetal bovine serum (FBS) at 37°C with 5% CO_2_ for three days and stained with India ink. Representative images are shown. (E) Quantification of the ratio of the capsule-to-cell body ratio from the experiment shown in (D). Violin plots show measurements from all analyzed cells, and dots represent the mean values from three independent biological replicates (*mob3*Δ-1, -2, and -3). (F) Histopathological analysis of lung tissue from WT- and *mob3*Δ-3-infected mice at 14-DPI. (G) Quantification of cell-body diameters and capsule thickness from histological images, shown as mean values from four independent animals per strain. Statistical significance was determined using one-way ANOVA with Dunnett’s multiple comparisons test (panels B, C, G), or an unpaired two-tailed Student’s *t* test (panel E). *In vitro* assays were performed using three biological replicates with three technical replicates each, whereas histopathological analyses included four animals per group.

Because interactions with phagocytes are critical determinants of *Cryptococcus* survival and dissemination during infection [45, 46], we next examined intracellular survival of wild type and *mob3*Δ strains within THP-1 macrophages. After 24 h of co-incubation with serum-opsonized *C. neoformans*, intracellular fungal burdens were significantly higher for *mob3*Δ strains than for wild type (Figure 6C). This enhanced recovery suggests that *mob3*Δ mutants have an increased capacity to persist within macrophages, potentially through increased resistance to phagosomal killing and/or altered intracellular proliferation [45, 47].

Given that cell size and capsule properties are known to influence both phagocytic interactions and endothelial transmigration, we next examined whether loss of *MOB3* alters fungal morphology under host-mimicking conditions. When grown in 10% serum at 37⁰C with 5% CO_2_, *mob3*Δ mutant cells exhibited reduced cell body size accompanied by noticeably enlarged capsules relative to wild type (Figure 6D). Quantification revealed a significantly increased capsule-to-cell-body ratio in *mob3*Δ mutants (Figure 6E), indicating that capsules occupied a greater proportion of the total cell size than in wild-type cells. These findings demonstrate that *MOB3* deletion promotes the formation of small-bodied, highly encapsulated cells under host-mimicking conditions.

To assess *mob3*Δ mutant cell morphology at the site of infection, mice were infected intranasally with wild type or *mob3*Δ strains, and lungs were collected at 14 days post infection for histopathological analysis (4 WT-infected animals and 12 animals infected with three independent *mob3*Δ strains) (Figure 6F). All animals showed marked lung infiltration and inflammation to a similar extent. However, *mob3*Δ infections were enriched for smaller fungal cells and contained fewer enlarged cells than wild-type infections. Quantification of fungal cells in randomly selected lung fields confirmed that all three independent *mob3*Δ strains had significantly reduced cell-body diameters relative to wild type whereas capsule size was not significantly different (Figure 6G). Consistent with these findings, histopathological analysis across multiple animals revealed smaller fungal cells and greater capsule-size variability in *mob3*Δ lung infections compared with wild type (Figure S8A). Notably, cells consistent with the previously described “seed cell” morphotype, which has been associated with enhanced dissemination [28], were more frequently observed in *mob3*Δ infections. In contrast, brain pathology was broadly comparable between WT- and *mob3*Δ-infected animals, with similar cryptococcoma formation and minimal surrounding host inflammation (Figure S8B). Together, these findings provide a mechanistic explanation for *mob3*Δ hypervirulence. Reduced cell size in the lung may facilitate escape from the respiratory tract, while enhanced BBB transmigration and increased intracellular survival within macrophages could further promote spread to the CNS and disease progression.

### Global phosphoproteomic profiling reveals component-specific signaling rewiring in STRIPAK mutants

Given the striking differences in virulence, morphology, and host adaptation among the STRIPAK mutants, we next performed global phosphoproteomic profiling to define the signaling pathways altered upon loss of individual STRIPAK components (Figure 7). We selected representative mutants occupying distinct functional positions within the STRIPAK complex (Figure S1), including the PP2A catalytic subunit (*pph22*Δ), the PP2A regulatory striatin subunit (*far8*Δ), the central scaffolding subunit (*far11*Δ), which exhibits severe phenotypes broadly representative of core STRIPAK dysfunction, and the Mob family signaling component (*mob3*Δ), which displays phenotypes opposite to those of core STRIPAK mutants. This strategy enabled assessment of both shared and subunit-specific signaling outputs downstream of STRIPAK disruption.

**Figure 7.**
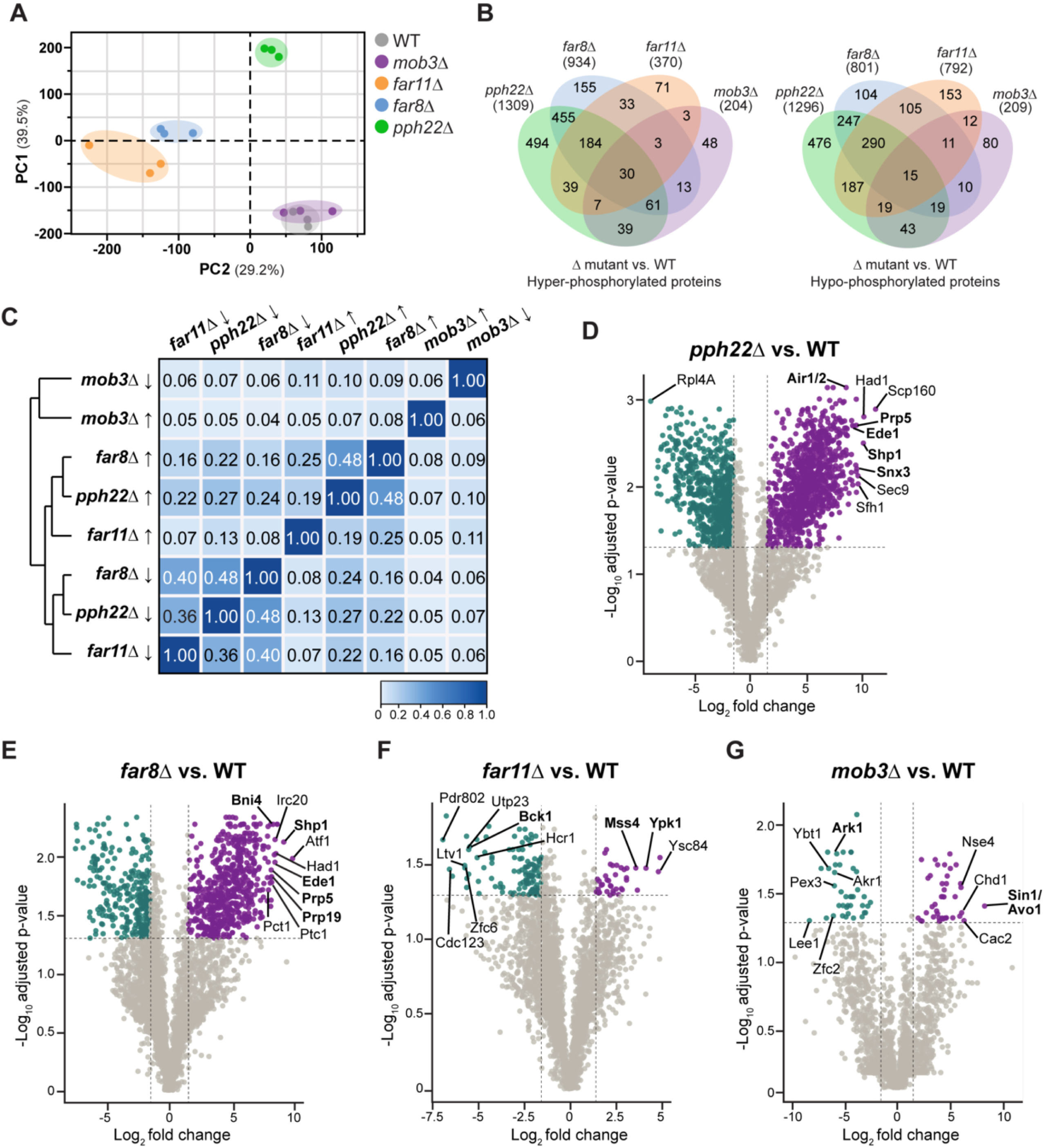
Global phospho-proteomic landscape of STRIPAK complex mutants. (A) Principal component analysis (PCA) of phosphoproteomic profiles from wild type and STRIPAK mutants shows that *pph22*Δ, *far8*Δ, and *far11*Δ samples cluster tightly together and are clearly separated from wild type along the major principal components, indicating extensive STRIPAK-dependent remodeling of the phosphoproteome. In contrast, *mob3*Δ samples cluster closer to wild type. (B) Venn diagrams illustrate the shared and unique phosphoproteins whose phosphorylation is significantly increased or decreased in *pph22*Δ, *far8*Δ, *far11*Δ, and *mob3*Δ relative to wild type. Comparison across mutants shows extensive overlap among the core STRIPAK components. The total number of identified unique phosphoproteins for each group is shown in parentheses. (C) A heatmap of pairwise Jaccard similarity scores (displayed within each cell) quantifies the degree of overlap among phospho-proteomic changes in the four STRIPAK mutants. (D-G) Volcano plots display significantly altered phosphoproteins in (D) *pph22*Δ, (E) *far8*Δ, (F) *far11*Δ, and (G) *mob3*Δ mutants compared to wild type. All phosphoproteins meeting reproducibility and high-confidence localization criteria across biological replicates are shown. Dashed lines indicate thresholds corresponding to |log_2_FC| ≥ 1.5 and -log10 adjusted p-values > 1.3, used to highlight strongly regulated candidates. Significantly up- (purple) and downregulated (teal) phosphoproteins are highlighted, and the ten most statistically significant hits in each mutant are labeled. Bolded proteins highlight representative phosphoproteins associated with the major signaling pathways discussed further in the text. The core STRIPAK mutants (*pph22*Δ, *far8*Δ, *far11*Δ) show widespread phosphorylation changes, whereas *mob3*Δ displays a comparatively smaller and distinct set of phosphorylation changes.

Phosphopeptides were enriched from biological triplicates of wild type and mutant strains and analyzed by LC–MS/MS. Differential phosphorylation was assessed using log_2_ - transformed intensities, and significantly regulated phosphopeptides were defined as those meeting both a fold-change threshold (≥1.5-fold) and a statistical cutoff (adjusted p-value < 0.1). Peptides passing these criteria were collapsed to unique proteins and classified as hyperphosphorylated (log₂FC > 1) or hypophosphorylated (log₂FC < –1) relative to wild type. Hyper- and hypophosphorylated protein sets were generated independently from the corresponding phosphopeptide datasets, such that proteins containing multiple regulated phosphosites could be represented in both sets. Principal component analysis revealed strong reproducibility within each genotype and clear separation of the core STRIPAK mutants *pph22*Δ, *far8*Δ, and *far11*Δ, all of which diverged sharply from wild type along the major principal components (Figure 7A). In contrast, *mob3*Δ mutant samples clustered tightly with wild type, consistent with the comparatively modest number of phosphorylation changes detected in this mutant.

To visualize shared and distinct STRIPAK-dependent phosphorylation events, we generated Venn diagrams of significantly hyper- and hypo-phosphorylated proteins across the mutants (Figure 7B, S3 Table). Mutants lacking the core STRIPAK subunits Pph22, Far8, or Far11 showed the strongest overlap, particularly among hypophosphorylated proteins, whereas *mob3*Δ exhibited far fewer regulated proteins and minimal overlap with the other mutants. To quantify these relationships, we computed pairwise Jaccard similarity scores across the eight protein sets (four mutants × up/down categories). The resulting heatmap (Figure 7C) confirmed that the downregulated sets of the *pph22*Δ, *far8*Δ, and *far11*Δ mutants were the most similar, followed by their upregulated sets. In contrast, *mob3*Δ up- and downregulated sets formed their own small, distinct clusters, reflecting the limited phosphoproteomic remodeling in this mutant. Together, these analyses demonstrate that loss of core STRIPAK components produces extensive and largely overlapping phosphoproteomic changes, whereas deletion of the *MOB3* gene results in a comparatively subtle and distinct phosphorylation profile, consistent with the phenotypic divergence between the hypervirulent *mob3*Δ mutant and the severely attenuated *pph22*Δ, *far8*Δ, and *far11*Δ mutant strains.

To identify the most strongly STRIPAK-regulated phosphoproteins in each mutant, we generated volcano plots highlighting proteins with both large effect sizes and strong statistical support (Figure 7D-G). The core STRIPAK mutants, *pph22*Δ, *far8*Δ, and *far11*Δ, each displayed hundreds of significantly hyper- or hypophosphorylated proteins, consistent with the extensive phosphoproteomic remodeling observed in these datasets. Several proteins involved in endocytic and membrane trafficking pathways (Ede1, Shp1, Snx3), pre-mRNA splicing and RNA processing (Prp5, Prp19, Air1/2), and cell polarity or cell wall organization (Bni4, Bck1) were among the most strongly affected phosphoproteins, suggesting broad STRIPAK-dependent control of signaling networks governing growth and cellular organization. Notably, *far11*Δ mutants also exhibited altered phosphorylation of proteins linked to TORC2 signaling and membrane homeostasis, including Mss4 and Ypk1. In contrast, *mob3*Δ mutants displayed a smaller set of high-confidence phosphorylation changes that included the actin-regulating kinase Ark1 and the TORC2 component Sin1/Avo1, suggesting that Mob3 influences a more restricted but distinct subset of STRIPAK-dependent signaling outputs.

To gain functional insight into the biological pathways most strongly affected by STRIPAK loss, we performed Gene Ontology (GO) enrichment analysis on the combined set of phosphoproteins that were significantly altered in at least two STRIPAK mutants. This filtering strategy ensured that the analysis reflected pathways commonly perturbed across the STRIPAK network rather than proteins uniquely affected in a single mutant background. The enriched GO terms (Figure 8, S4 Table) grouped into several major functional modules, including signaling and membrane dynamics, cell-cycle and cytoskeletal regulation, protein modification and post-translational control, chromatin and transcriptional regulation, and metabolic and stress-response pathways. Many of these enriched categories are consistent with the phenotypes observed in STRIPAK mutants, including defects in cell-cycle progression, genome stability, cell morphology, stress adaptation, and capsule regulation. In particular, enrichment of pathways related to small GTPase signaling, membrane dynamics, mitotic checkpoint regulation, DNA replication, and cytoskeletal organization suggests that STRIPAK coordinates signaling networks that govern both cellular architecture and genome maintenance. Additional enrichment of chromatin, RNA-metabolism, and stress-response pathways points to broader roles for STRIPAK in regulating adaptive cellular programs.

**Figure 8.**
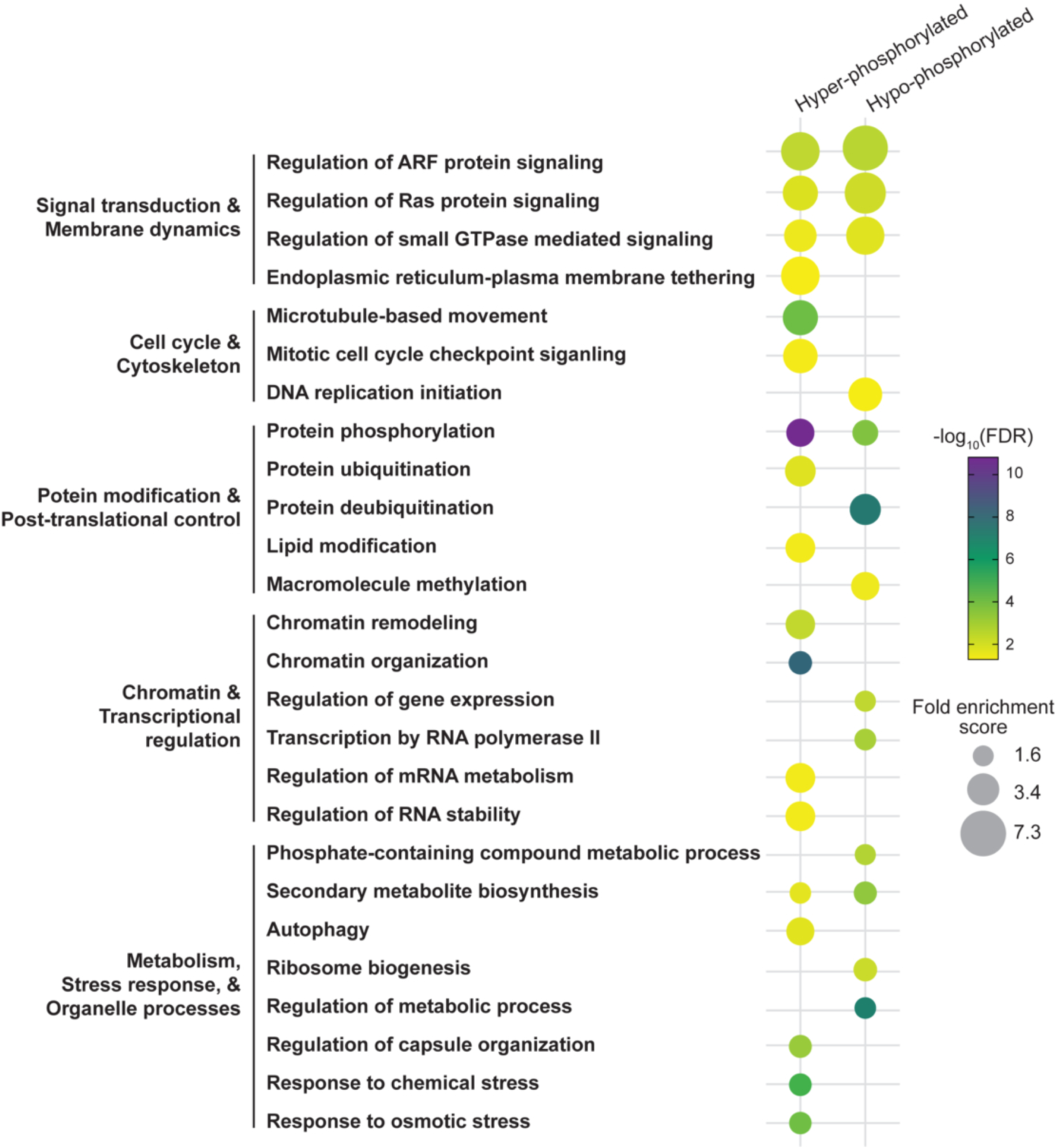
Gene Ontology (GO) term enrichment analysis of STRIPAK-dependent phosphoproteome changes. GO biological process terms significantly enriched among hyper-phosphorylated (left) and hypo-phosphorylated (right) proteins across STRIPAK mutants are visualized as a bubble plot. GO enrichment was performed using the combined set of phosphoproteins that met significance cutoffs in at least two STRIPAK mutants, ensuring that the analysis focused on commonly affected pathways rather than proteins altered in only a single mutant background. GO terms are grouped into major functional categories, including signaling and membrane dynamics, cell-cycle and cytoskeletal processes, protein modification, chromatin and transcriptional regulation, and metabolic and stress response pathways. Bubble size reflects fold-enrichment score, and bubble color indicates statistical significance (–log_10_ FDR). Enriched GO terms span diverse biological processes, indicating that STRIPAK-dependent phosphorylation changes affect multiple interconnected cellular pathways.

To place STRIPAK-dependent phosphorylation changes into a systems-level context, we next examined the protein-protein interaction architecture of differentially phosphorylated proteins using STRING [48] with a high-confidence interaction threshold (score ≥ 0.70) restricted to experimental and curated database evidence (Figure 9). The resulting network resolved into multiple discrete functional modules, highlighting the breadth of STRIPAK-regulated signaling. A large cluster of Ras/GTPase- and MAPK-associated proteins included core cell wall integrity and stress-activated kinases (Pkc1, Bck1, Ste7, Mkk2, Hog1), as well as nutrient- and growth-responsive kinases (Pka1/Pkr1, Cac1, Sch9, Cbk1, Pak1). This module also included polarity and cytoskeleton organizers Bem1, Iqg1, Rga3, Spa2, and Bni1, suggesting coordinated STRIPAK control over signaling cascades that couple environmental sensing with cytoskeletal remodeling and morphogenesis. Together, these proteins comprise major pathways controlling cell-wall integrity, osmotic stress adaptation, nutrient sensing, and polarized growth, consistent with the morphological and stress-response phenotypes observed in STRIPAK mutants.

**Figure 9.**
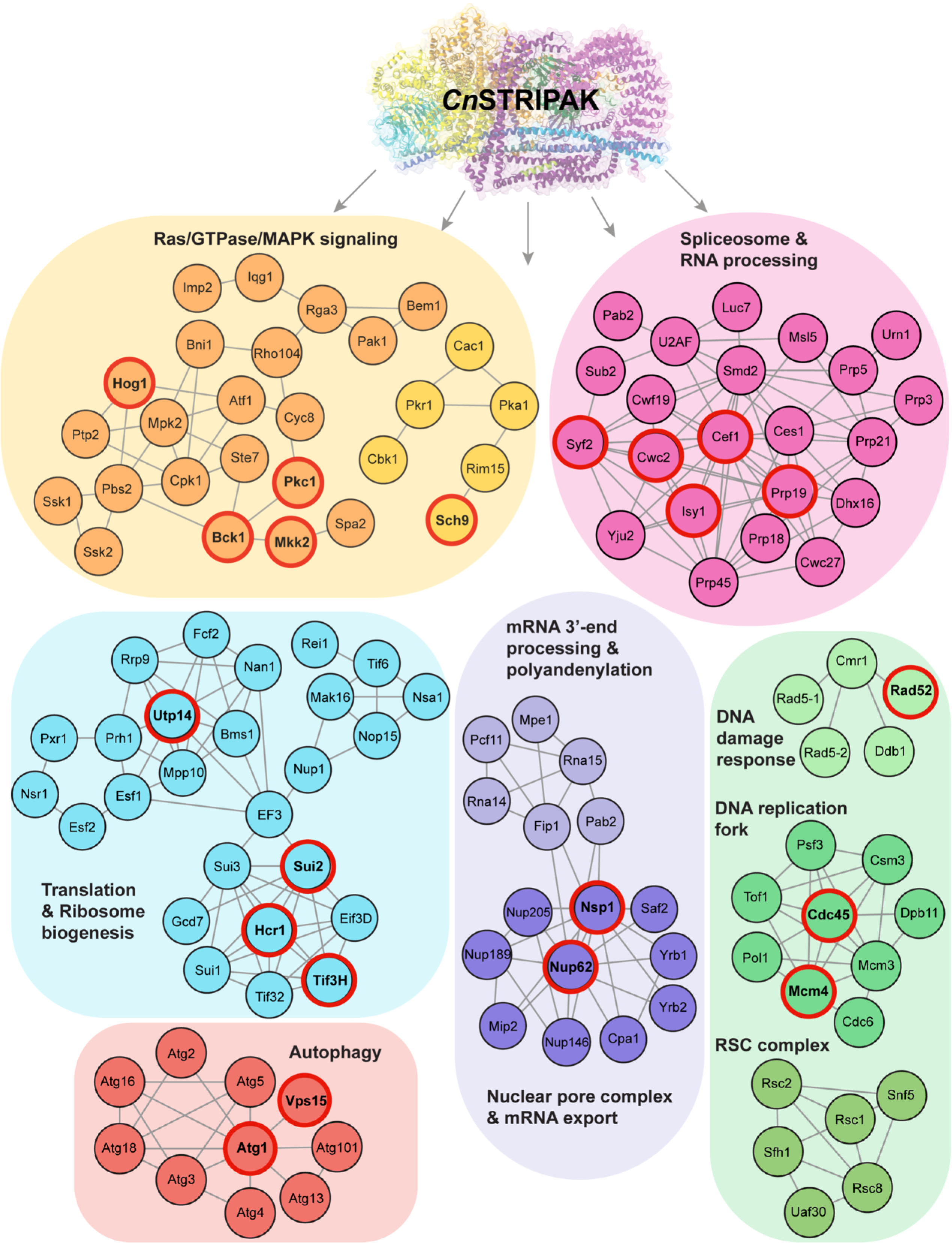
Predicted STRIPAK-dependent signaling networks among differentially phosphorylated proteins. Proteins with significant differential phosphorylation compared to wild type were analyzed using STRING with a high-confidence interaction threshold (score ≥ 0.70) and evidence restricted to experimental data and curated databases. The resulting interaction map resolves into distinct functional modules representing Ras/GTPase/MAPK signaling, RNA processing and splicing, chromatin remodeling, DNA replication and repair, translation and ribosome biogenesis, and autophagy. Nodes are colored by functional category, and edges represent high-confidence protein-protein associations (STRING score ≥ 0.70). Selected representative proteins are outlined in red to illustrate key components of the identified functional modules.

A second major module comprised spliceosome and RNA-processing machinery, including components of the conserved NTC/Prp19 complex together with additional factors involved in pre-mRNA splicing and transcript maturation. This network was closely connected to proteins involved in nuclear transport and mRNA export, including nucleoporins, Ran-associated transport factors, and components of the 3′-end processing and polyadenylation machinery. These interconnected modules suggest that STRIPAK-dependent phosphorylation influences multiple stages of RNA metabolism, spanning transcript processing, nuclear export, and post-transcriptional gene regulation.

Additional functional groups included an autophagy module comprising the Atg1-Atg13-Atg101 initiation complex together with downstream membrane-expansion components; a DNA replication fork module containing CMG (Cdc45-MCM-GINS) helicase components, replication initiation factors, and checkpoint proteins; and a DNA damage-response network. The enrichment of replication-initiation, fork-protection, and DNA damage-response factors is consistent with the genome instability, copy-number variation, and abnormal DNA-content profiles observed in STRIPAK mutants.

We also identified the RSC chromatin-remodeling module and an extensive network associated with translation initiation and ribosome biogenesis, with elongation factor 3 (EF3) linking multiple submodules. These networks suggest that STRIPAK-dependent signaling extends beyond classical kinase signaling pathways to influence chromatin organization, gene expression, and core biosynthetic processes. Representative proteins are outlined in red to highlight key components within each functional module, including signaling kinases as well as proteins involved in RNA splicing, nuclear transport, DNA replication, autophagy, and ribosome biogenesis (Figure 9). Together, these proteins represent candidate downstream effectors of STRIPAK-dependent signaling.

Among the phosphoproteins comprising the shared hyperphosphorylated protein set were several proteins associated with TORC2 signaling and lipid homeostasis, including Efr3, Mss4, Pkh2, and Ypk1, together with proteins involved in endocytosis and actin cytoskeleton organization, including Cin1, Sla2, and Ark1 (Figure 10A). In fungi, TORC2 signaling regulates membrane and lipid homeostasis, cell-wall integrity, actin polarization, and cellular morphogenesis, while endocytosis pathways coordinate membrane remodeling and protein trafficking [49-51]. The identification of multiple proteins within these interconnected pathways suggests that dysregulation of TORC2-associated signaling and membrane trafficking may represent a common downstream consequence of STRIPAK disruption.

**Figure 10.**
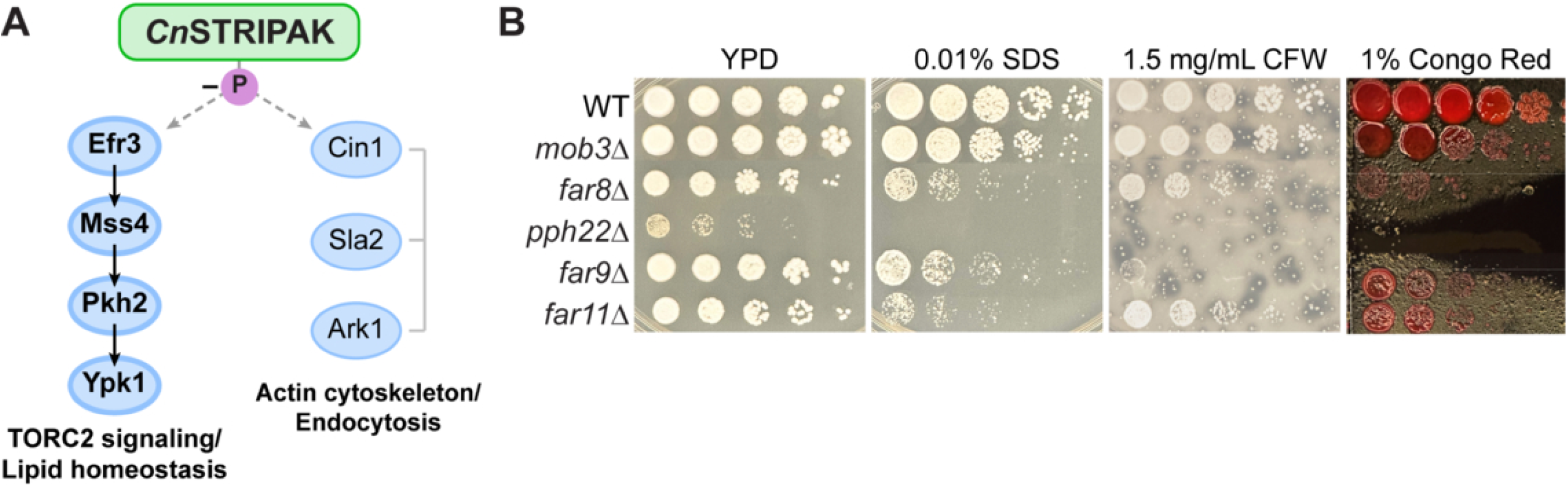
Shared STRIPAK-dependent phosphoproteins implicate TORC2 signaling and membrane homeostasis pathways. (A) Selected proteins from the set of 30 phosphoproteins that were significantly hyperphosphorylated across all STRIPAK mutants (center overlap in Figure 7B). Proteins were grouped according to their predicted roles in the TORC2 signaling axis, lipid homeostasis, actin cytoskeleton organization, and endocytosis. Arrows denote established signaling relationships within the TORC2 pathway and are included for conceptual illustration. (B) Serial dilution spotting assays of WT and STRIPAK mutant strains under conditions that perturb membrane and cell-wall homeostasis. Cells were serially diluted and spotted onto YPD, YPD supplemented with 0.01% SDS, 1.5 mg/mL calcofluor white (CFW), or 1% Congo red and incubated at 30°C. Images were captured after 2 days of incubation.

To determine whether STRIPAK mutants exhibit phenotypes consistent with altered TORC2 pathway activity, we examined growth under conditions that perturb membrane and cell-wall homeostasis. Multiple STRIPAK mutants displayed increased sensitivity to SDS, calcofluor white (CFW), and Congo red (Figure 10B). Sensitivity was most severe in *pph22*Δ and *far11*Δ mutants, whereas *far8*Δ and *far9*Δ also exhibited pronounced growth defects under these conditions. In contrast, *mob3*Δ displayed comparatively modest defects on SDS and CFW but remained sensitive to Congo red. Together, these findings provide functional support for the phosphoproteomic data and implicate TORC2- and membrane homeostasis-associated pathways as candidate downstream targets of STRIPAK signaling.

Together, the phosphoproteomic, enrichment, and network analyses demonstrate that STRIPAK-dependent phosphorylation changes converge on interconnected regulatory hubs controlling stress responses, morphogenesis, genome maintenance, RNA metabolism, autophagy, protein synthesis, and membrane homeostasis. These findings provide a mechanistic framework linking STRIPAK dysfunction to the diverse developmental, morphological, and virulence-associated phenotypes observed throughout this study and identify TORC2-associated signaling as a candidate pathway underlying STRIPAK-dependent regulation of fungal morphogenesis and host adaptation.

## Discussion

In this study, we establish the STRIPAK complex as a central signaling hub that coordinates genome stability, cellular morphology, stress adaptation, and virulence in *C. neoformans*. Building on our previous work demonstrating roles for STRIPAK components in growth, development, and pathogenicity [33, 34], we now extend these findings by integrating phenotypic, genomic, *in vivo* infection, and systems-level phosphoproteomic analyses across multiple STRIPAK subunits. Together, these findings demonstrate that loss of the PP2A catalytic subunit Pph22, the regulatory striatin subunit Far8, or the adaptor proteins Far9 and Far11 disrupts core STRIPAK signaling required to integrate cell-cycle progression, genome maintenance, and stress-responsive growth in *C. neoformans*, underscoring the fundamental role of STRIPAK-mediated phosphatase signaling in fungal cellular homeostasis [52]. Our data also reveal subunit-specific outputs, most notably for Mob3, whose loss produces phenotypes opposite to those observed for other STRIPAK components and confers enhanced virulence-associated traits during host infection. Global phosphoproteomic profiling provides a mechanistic framework for these observations, uncovering extensive overlap in phosphorylation networks regulated by core STRIPAK components. Together, the convergence of systems-level signaling data with cellular and infection phenotypes highlights STRIPAK as an organizing platform that integrates multiple regulatory pathways, enabling *C. neoformans* to balance genome stability, morphological plasticity, and environmental adaptation during infection.

Disrupted STRIPAK signaling results in profound genome instability in *C. neoformans*, manifested as widespread whole-chromosome aneuploidy and progression toward polyploidization. Whole-genome sequencing and flow cytometry of *far9*Δ and *far11*Δ mutant strains revealed heterogeneous populations with altered chromosome copy number and increased DNA content consistent with progression toward polyploidization. Consistent with these phenotypes, our phosphoproteomic analyses suggest STRIPAK-dependent regulation of pathways central to chromosome segregation and genome stability, including cell-cycle regulators, microtubule-associated proteins, DNA replication factors, and chromatin-remodeling complexes [53, 54]. In addition, several phospho-regulated proteins identified across core STRIPAK mutants are associated with membrane signaling pathways that coordinate cellular growth with stress adaptation, suggesting that STRIPAK may influence genome stability through broader signaling networks that integrate cell-cycle progression with environmental responses. This role for STRIPAK in genome maintenance is consistent with observations in other fungi, where STRIPAK components regulate septation, nuclear division, and mitotic progression [11, 55-58]. In mammalian systems, *Drosophila,* and *C. elegans*, STRIPAK-PP2A complexes regulate centrosome function, spindle orientation, and microtubule organization [59-63]. Our findings extend these conserved functions to a human fungal pathogen and demonstrate that STRIPAK-mediated phosphatase signaling is essential for buffering genome instability in the face of environmental and host-imposed stresses. In *C. neoformans*, failure to maintain this coordination results in genome instability that may initially impair growth but also provides a basis for adaptive evolution during infection. Together, these data suggest that STRIPAK stabilizes the genome not through a single linear pathway, but by coordinating phosphorylation states across multiple interdependent processes that ensure accurate DNA replication, spindle function, and chromatin organization.

Disruption of core STRIPAK components also results in pronounced morphological heterogeneity, linking genome instability to altered cell size and cell-cycle control. *far9*Δ and *far11*Δ mutants produced enlarged cells, exhibited defects consistent with impaired cytokinesis, and formed titan cell-like morphologies even under non-inducing conditions [40]. These phenotypes coincided with stable chromosome copy number alterations and elevated DNA-content states that persisted across growth conditions, indicating that defects in genome maintenance is a sustained consequence of STRIPAK dysfunction rather than a transient stress response. This contrasts with stress-induced polyploidization in *Cryptococcus*, where unchecked DNA replication coupled with cell-cycle arrest can drive a temporary expansion of polyploid cells [64]. One plausible explanation is that STRIPAK-dependent phosphorylation normally coordinates cell-cycle progression with cytoskeletal organization, including kinetochore function, centrosome dynamics, and actin remodeling [65], such that loss of this coordination uncouples nuclear division from cytokinesis and promotes cell enlargement [66]. Notably, these findings extend observations from multicellular eukaryotes, where STRIPAK dysregulation has been linked to tumorigenesis, to a fungal pathogen context [65, 67, 68].

Our phosphoproteomic data further suggest a functional connection between STRIPAK and TORC2 signaling. Multiple regulators and effectors of TORC2 signaling, including Efr3, Mss4, Pkh2, and Ypk1, were identified among the differentially phosphorylated phosphoproteins in STRIPAK mutants, supporting a role for STRIPAK in regulating plasma membrane homeostasis, endocytosis, and actin-dependent morphogenesis. Efr3 and Mss4 function upstream of TORC2 by promoting phosphatidylinositol-4,5-bisphosphate production at the plasma membrane, while Pkh2 and Ypk1 are central regulators of membrane stress responses, polarized growth, and cytoskeletal organization. Given established roles for TORC2-Ypk1 signaling in coordinating membrane growth with cytokinesis and cell expansion, disruption of this regulatory axis provides a potential mechanistic explanation for the enlarged-cell phenotypes, cytokinesis defects, and morphological heterogeneity observed in *far9*Δ and *far11*Δ mutants. Together, these findings support a model in which STRIPAK coordinates cell-cycle progression and morphogenesis, at least in part, through modulation of TORC2-dependent signaling pathways.

During infection, *far9*Δ mutants displayed striking morphological diversity *in vivo*, including titan cells, microcells, and pseudohyphal forms within the same host tissues. This heterogeneity mirrors established adaptive strategies in *C. neoformans*, in which enlarged titan cells can evade phagocytosis and immune clearance, while smaller cell types (seed cells) are associated with enhanced dissemination [28, 46, 69]. Despite severe stress sensitivity *in vitro*, *far9*Δ mutant cells disseminated to the brain by day 14 post-infection in all animals but did not cause overt disease until weeks later, revealing a paradoxical virulence phenotype characterized by delayed yet ultimately fatal infection. Notably, isolates recovered during infection acquired additional chromosome copy number changes without displaying a clear fitness advantage under *in vitro* conditions, suggesting that genomic alterations may confer a context-dependent benefit within the host. One possibility is that *far9*Δ mutant cells initially persist in a low-replicative or quiescent state within the CNS, delaying immune activation or inflammatory pathology. In support of this model, recent work has shown that *Cryptococcus* can rapidly infiltrate the brain while eliciting a markedly delayed microglial and leukocyte response, resulting in an extended window for establishment of infection prior to symptomatic disease [70]. Together, these findings suggest that disrupted STRIPAK signaling may enable persistence and delayed disease progression under physiological host conditions [71].

In contrast to core STRIPAK components whose loss primarily disrupts development and stress tolerance, Mob3 emerged as a functionally divergent regulator with effects that are most pronounced in the host environment. Despite relatively modest phosphoproteomic changes under standard *in vitro* conditions, *mob3*Δ mutants exhibited striking virulence-associated phenotypes, including enhanced transmigration across an *in vitro* blood-brain barrier and survival within macrophages, increased formation of microcells or “seed” cells, and disproportionate capsule expansion relative to cell body size. Previous work has shown that macrophage phagocytosis of *C. neoformans* is strongly influenced by capsule size, with smaller-capsule cells preferentially internalized during early infection, while subsequent capsule enlargement rapidly limits further uptake and can enhance survival within the phagosome [47, 72]. In this context, the increased capsule-to-cell-body ratio observed in *mob3*Δ strains is consistent with a strategy that promotes persistence within the macrophage, providing a plausible cellular basis for the enhanced recovery of viable *mob3*Δ mutant cells following macrophage co-incubation. Together, these traits are closely linked to dissemination and CNS invasion, suggesting the consequences of *MOB3* deletion are selectively engaged during infection. Rather than broadly regulating phosphorylation, our data suggest that Mob3 directs STRIPAK activity toward distinct substrates or cellular contexts that are not apparent under routine laboratory conditions, thereby restricting the expression of specific virulence-associated traits.

This model is consistent with findings in other systems, where Mob family proteins act as scaffolds and allosteric regulators that tune kinase signaling outputs, including within STRIPAK-associated modules [63]. In metazoans, STRIPAK antagonizes Hippo signaling, and Mob proteins direct pathway specificity by coordinating kinase activation states [73]. Together, these observations support a model in which Mob3 serves as a context-dependent STRIPAK regulator that shapes infection-relevant cellular behaviors. These findings underscore the importance of STRIPAK, and Mob3 in particular, as a modular signaling node whose subunit-specific regulation is selectively engaged in the host to influence pathogenic potential.

Our phosphoproteomic and network analyses position STRIPAK as a central organizer of *Cryptococcus* cell signaling. Across core STRIPAK mutants, we observed extensive overlap in differentially phosphorylated proteins and highly interconnected interaction networks, implicating shared regulation of pathways controlling kinase signaling, cell-cycle progression, cytoskeletal dynamics, chromatin organization, RNA processing, autophagy, and ribosome biogenesis. STRING network analysis further revealed that many STRIPAK-regulated phosphoproteins cluster into coherent functional modules and include multiple kinases, supporting the model that PP2A-containing STRIPAK complexes can act as kinase-directed phosphatases that tune signaling outputs across pathways [5]. The convergence of shared kinase targets across multiple STRIPAK mutants highlights the role of STRIPAK as a signaling hub that integrates phosphorylation networks to maintain cellular homeostasis [17]. Notably, multiple components of the TORC2 signaling pathway and its upstream regulators were identified among STRIPAK-regulated phosphoproteins. Together with the morphological and cytokinetic defects observed in core STRIPAK mutants, these findings support a model in which STRIPAK modulates TORC2-dependent signaling through phosphorylation changes affecting multiple upstream regulators and downstream effectors, including Efr3, Mss4, Pkh2, and Ypk1. Given the established roles of this pathway in plasma membrane homeostasis, endocytosis, actin organization, and stress adaptation, STRIPAK-dependent regulation of TORC2 signaling provides a mechanistic framework linking the phosphoproteomic changes observed here to the genome instability, morphogenetic defects, and stress-sensitive phenotypes of STRIPAK mutants.

These systems-level insights help explain how perturbation of STRIPAK can simultaneously impact genome stability, morphology, stress adaptation, and virulence, and why disruption of individual subunits produces both shared and divergent phenotypic outcomes. Our findings support a model in which STRIPAK functions as a central phosphatase signaling hub that coordinates multiple cellular processes through shared phosphorylation networks and, in particular, through modulation of TORC2-associated signaling pathways involved in membrane homeostasis, endocytosis, and morphogenesis. By coordinating signaling across diverse cellular processes, STRIPAK provides a framework for rapid phenotypic adaptation during infection, while subunit-specific regulation enables context-dependent rewiring of signaling outputs in the host. Given the broad conservation of STRIPAK components but the presence of fungal-specific protein features and complex organization, together with the subunit-specific virulence phenotypes uncovered here, STRIPAK represents a potential target for antifungal strategies aimed at disrupting regulatory hubs rather than individual pathways, provided that uniquely fungal aspects of the complex can be exploited to minimize host toxicity [74, 75]. Future work will focus on defining STRIPAK-dependent phosphorylation dynamics under host-relevant conditions and experimentally validating high-confidence substrate candidates identified by our network analyses, thereby providing deeper insight into how conserved phosphatase signaling networks are repurposed to drive fungal pathogenicity.

## Materials and methods

### Strains, media, and growth conditions

*Cryptococcus* strains used in this study are listed in Table S1, which were prepared for long-term storage as 20% glycerol stocks at -80°C. Fresh cultures for each experiment were revived on YPD agar medium (1% yeast extract, 2% Bacto Peptone, 2% dextrose). Transformed fungal strains were selected on YPD medium supplemented with 100 μg/mL nourseothricin (NAT) and/or 200 μg/mL neomycin (G418). For plate assays, strains were grown on Murashige and Skoog (MS) (Sigma-Aldrich M5519), YNB (0.67% yeast nitrogen base with ammonium sulfate and without amino acids, 2% dextrose), Niger seed (7% Niger seed, 0.1% dextrose), or L-3,4-dihydroxyphenylalanine (L-DOPA) (7.6 mM L-asparagine monohydrate, 5.6 mM glucose, 22 mM KH_2_PO_4_, 1 mM MgSO_4_.7H_2_O, 0.5 mM L-DOPA, 0.3 μM thiamine-HCl, 20 nM biotin, pH= 5.6). All plate media were prepared with 2% Bacto agar. L-DOPA plates were supplemented with 10 μM CuSO_4_ or 10 μM of the copper chelator bathocuproine disulfonate (BCS) to assess copper dependency.

To analyze stress response phenotypes, YPD medium was supplemented with 1 μg/mL FK506, 100 μg/mL cyclosporine A, 1 M sorbitol, 1.5 M NaCl, 1.5 M KCl, 0.01% sodium dodecyl sulfate (SDS), 3 mg/mL calcofluor white (CFW), 0.5 mg/mL caffeine, or 1% Congo red. Fluconazole Etests were performed using 0.016–256 μg/mL MIC test strips (Liofilchem 921470). For capsule analysis, strains were incubated for 3 days in RPMI media or 10% fetal bovine serum [heat-inactivated and diluted in phosphate-buffered saline (PBS)] at 30°C, 37°C, or 37°C with 5% CO_2_, followed by counter-staining with India ink. For serial dilution assays, fresh cell cultures were diluted to equal OD_600_, serially diluted 5-fold, and spotted onto plates for the indicated conditions. Plates were incubated for 2 to 7 days and photographed daily.

### Construction of mutant strains

Deletion mutant strains were generated in the *C. neoformans* H99α or KN99**a** backgrounds. The deletion alleles were constructed by fusing approximately 1 kbp 5’ and 3’ homologous arms, amplified from H99α genomic DNA, of the target gene to nourseothricin acetyltransferase (NAT) or neomycin resistance (NEO) gene expression cassettes. Homologous arms were assembled with the drug resistance marker with overlap PCR [76, 77]. The *far9*Δ*::NAT* and *far11*Δ*::NAT* mutants were generated via biolistic transformation, according to previously described methods [78, 79]. Complemented strains *far9*Δ*::GFP-FAR9-NEO* and *far11*Δ*::FAR11-NEO* were constructed via CRISPR-Cas9-directed mutagenesis [80, 81]. For the *far9*Δ*::GFP-FAR9-NEO* strain, an N-terminal GFP-tagged *FAR9* allele together with a neomycin resistance cassette (NEO^R^) was integrated upstream of the *far9*Δ::*NAT* deletion cassette. For the *far11*Δ::*FAR11-NEO* strain, the *FAR11* allele and NEO^R^ cassette were reintroduced at the endogenous *FAR11* locus. Stable transformants from YPD+NAT or YPD+NEO selection plates were screened by diagnostic PCR to confirm integration of the desired allele at the endogenous locus. Positive transformants were further confirmed via Illumina whole-genome sequencing. Primers used in this study are listed in Table S2.

### Reverse transcriptase quantitative PCR (RT-qPCR)

Total RNA was extracted from mid-logarithmic phase cultures using the RNeasy Plant Mini Kit (Qiagen) with on-column DNase treatment according to the manufacturer’s instructions. cDNA was synthesized from 1 μg of DNase-treated RNA using the iScript cDNA Synthesis Kit (Bio-Rad). Quantitative PCR reactions were performed using PowerUp SYBR Green Master Mix (Applied Biosystems) on a QuantStudio 3 system (Thermo Fischer Scientific). Relative transcript abundance was normalized to the housekeeping gene *GAPDH* and calculated using the 2^^-ΔΔCt^ method [82]. Statistical analyses were performed on ΔCt values. Each experiment included three biological replicates, with two technical replicates per biological replicate.

### Flow cytometry

Ploidy of *Cryptococcus* strains was analyzed via fluorescence-activated cell sorting (FACS) as previously described, with minor modifications [83, 84]. Wild-type and mutant strains were grown overnight in YPD or 10% FBS medium at 30°C or 37°C, harvested, and washed with PBS. Cells were then fixed in 70% ethanol for 16 hours at 4°C. Fixed cells were washed with 1 mL NS buffer (10 mM Tris-HCl, pH=7.5; 250 mM sucrose; 1 mM EDTA, pH= 8.0; 1 mM MgCl_2_; 0.1 mM CaCl_2_; 0.1 mM ZnCl_2_; 0.4 mM phenylmethylsulphonyl fluoride; and 7 mM β-mercaptoethanol) and treated with RNase A (0.5 mg/mL). Cells were stained with propidium iodide (10 μg/mL) in NS buffer for 2 hours in the dark. Immediately prior to flow cytometry, 50 μL of stained cells were diluted into 50 mM Tris-HCl (pH7.5) and sonicated to minimize cell clumping. Samples were submitted to the Duke Cancer Institute Flow Cytometry Shared Resource for analysis. Fluorescence was measured using a BD FACSCanto flow cytometer and analyzed with FlowJo software. Doublets and cell aggregates were excluded using pulse-width discrimination (PI-W vs. PI-A), and only singlet events were included in subsequent DNA-content analyses. Haploid (H99α) and diploid (KN99α/KN99**a**) controls were included in each experiment for DNA-content calibration. Approximately 15,000 events were analyzed per sample.

### Microscopy and quantification

Brightfield, differential interference contrast (DIC), and fluorescence microscopy images were visualized with an AxioScop 2 fluorescence microscope equipped with an AxioCam MRm digital camera (Zeiss, Germany). Consistent exposure times were used for all images collected within each experiment. Cell body diameters were measured in ImageJ/Fiji and polysaccharide capsule thickness was calculated using the Quantitative Capture Analysis program [85]. At least 60 cells were measured per biological replicate. Statistical differences in cell body size and capsule thickness between test groups were assessed by one-way ANOVA, followed by Dunnett’s multiple comparisons test (for comparisons to the wild-type control) or Tukey’s test (for all pairwise comparisons). Statistical differences in capsule thickness-to-cell body radius ratios were evaluated using an unpaired t-test with Welch’s correction to account for unequal variances.

For *in vivo* analyses of fungal cell morphology within host tissue, lung homogenates collected from mice at 14 days post-infection were washed twice in PBS and stained with 10 μg/mL Calcofluor white (CFW) to label fungal cell walls and distinguish them from surrounding host tissue. Stained samples were incubated in the dark for 15 minutes, washed once with PBS, and imaged immediately. Fungal chitin was visualized using a UV fluorescence filter to detect the characteristic blue Calcofluor signal.

### Mating and sexual reproduction analysis

To assess mating between *Cryptococcus* strains, *MAT***a** and *MAT*α strains were grown overnight in YPD liquid media at 30°C, mixed, and spotted onto MS plates for incubation at room temperature in the dark. For the wild-type control cross, KN99**a** and H99α cells were mixed in equal amounts before spotting. For crosses involving *far9*Δ and *far11*Δ mutant strains, which exhibit significant growth defects at room temperature (25°C), mutant cells were first spotted alone on MS plates and allowed to pre-grow for two days in the absence of the wild-type partner. After pre-incubation, opposite mating type wild-type cells were spotted on top of the mutant cells. Mating plates were monitored and imaged for six weeks to document filamentation, basidia formation, and basidiospore production. Sexual development phenotypes from mutant × wild-type crosses were evaluated across at least 6 biological replicates.

### Whole-genome sequencing

Genomic DNA for whole-genome sequencing was extracted from YPD-grown cells using the MasterPure Yeast DNA Purification Kit (LGC Biosearch Technologies, MPY80200). Precipitated DNA was dissolved in 1x TE buffer (100 mM Tris-HCl, 10 mM EDTA, pH= 8.0) and concentration and purity were estimated using Qubit and Nanodrop, respectively. Illumina sequencing was performed at the Duke Sequencing and Genomic Technologies core facility with Novaseq 6000 providing 150 bp paired-end reads. Trimmed Illumina sequencing reads were mapped to the *C. neoformans* H99α reference genome using BWA-MEM2 through the Galaxy web server (https://usegalaxy.org/). The resulting BAM files were imported into Geneious for SNP calling using the Geneious default mapper with five iterations, a 0.9 variant frequency threshold, and a minimum coverage of 90x per variant. To assess chromosome copy number variation, BAM files were converted to bedgraph format and genome-wide read-depth coverage was calculated in 500-bp bins. For each sample, coverage values were normalized to the median genome-wide coverage and plotted across concatenated chromosomes in R (v4.5.3) using ggplot2. Coverage plots were generated using a modified version of the publicly available pipeline described by Lehmann et al., 2026 [86].

### Murine infection model and histopathology

*C. neoformans* inoculum was prepared by culturing fresh cells in YPD at 30°C for approximately 8 generations. Cells were collected by centrifugation and washed twice with sterile PBS. The cell density was determined with a hemacytometer and was adjusted to 4 x 10^6^/mL in PBS. Five- to six-week-old A/J mice (Jackson Laboratory, USA) were utilized for the murine intranasal infection model (n=14 for each group, 7 male and 7 female). Mice were anesthetized with isoflurane and infected by intranasal instillation of a 25 μL inoculum of 10^5^ cells. Animal survival was monitored daily and euthanasia was performed via CO_2_ exposure upon reaching humane endpoints, including weight loss greater than 20%, hunched appearance, or reduced grooming and mobility. For fungal burden analysis, two male and two female mice from each group were randomly selected and euthanized at 14 days post-infection. Brains and lungs were dissected and homogenized in 1 mL PBS using a bead beater. Organ homogenates were serially diluted and plated onto YPD medium supplemented with chloramphenicol (30 μg/mL) and ampicillin (100 μg/mL) to recover fungal colonies. Individual colonies recovered from lung or brain homogenates were re-streaked for single colonies prior to whole-genome sequencing and phenotypic characterization. Mouse survival was plotted using Kaplan-Meier curves and analyzed by the log-rank test. Fungal burden data were analyzed using one-way ANOVA with Dunnett’s multiple comparisons test.

In an independent experiment, mice were infected as described above, and four animals per group were selected for histopathological analysis either at 14 days post-infection (for *mob3*Δ mutants and the KN99**a** wild-type control) or upon reaching humane endpoints (approximately day 45 for *far9*Δ mutant-infected mice and day 20 for wild-type H99α). Brain, lungs, heart, spine, ribs, and associated lymph nodes were dissected and preserved in 10% neutral-buffered formalin for histological processing at the Duke Research Animal Pathology core facility.

### Blood-brain barrier transmigration assay

To analyze the ability of *Cryptococcus* strains to cross a blood–brain barrier (BBB) model, a transwell assay was performed as previously described [43, 87]. Human brain microvascular endothelial cells (hCMEC/D3) were grown to approximately 80% confluency and seeded at a density of 5 × 10⁴ cells onto collagen-coated transwell inserts containing 8 µm pore size permeable membranes. Cells were cultured in EndoGRO medium (SCME004, Sigma-Aldrich, USA) at 37⁰C with 5% CO_2_ for 24 h, after which the medium was replaced with EndoGRO media supplemented with 2.5% human serum, and monolayers were allowed to mature for an additional four days. One day prior to fungal inoculation, fresh medium supplemented with 1.25% human serum was added to promote endothelial attachment and tight junction formation. Integrity of the endothelial monolayer was assessed by measuring transendothelial electrical resistance (TEER) using an Epithelial Volt per Ohm Meter (EVOM3 device, World Precision Instruments, USA) and only inserts exhibiting stable resistance values were used for transmigration assays. Wild-type and mutant *Cryptococcus* strains were grown overnight, washed three times with PBS, and adjusted to a concentration of 5 × 10^5^ cells in 100 µL PBS using a hemocytometer. Fungal cells were added to the apical chamber of hCMEC/D3-seeded transwells and incubated for 24 hours at 37⁰C with 5% CO_2_. Following incubation, samples were collected from both the apical chamber and the basolateral compartment to recover fungal cells that had transmigrated across the endothelial monolayer. The BBB crossing ratio for each mutant strain was calculated and normalized to the wild-type control to determine relative transmigration efficiency. Statistical differences between wild-type and mutant groups were assessed using one-way ANOVA followed by Dunnett’s multiple comparisons test.

### Intracellular survival assay of *C. neoformans* within macrophages

Preparation of macrophage and *C. neoformans* cultures, infection of macrophages, and assessment of fungal survival were performed as previously described, with minor modifications [88-91]. THP-1 cells were seeded at 600 μL of 3.33 x 10^5^ cells/mL in 24-well tissue culture plates and differentiated in RPMI 1640 medium (ATCC modification; Gibco) supplemented with 10% FBS with 32 nM phorbol 12-myristate 13-acetate (PMA) for 48 h at 37⁰C with 5% CO_2_, followed by a 24 h recovery period in media without PMA. *C. neoformans* cells from overnight cultures were washed twice with PBS, adjusted to 1 x 10^8^ cells/mL, and opsonized in RPMI 1640 medium containing 40% FBS for 30 min at 37⁰C. Opsonized fungi were washed in PBS, resuspended in RPMI 1640, and added to differentiated THP-1 monolayers at a multiplicity of infection (MOI) of 5. Co-cultures were incubated at 37⁰C with 5% CO_2_ for 24 h.

Following co-incubation, the culture medium was aspirated, and wells were washed twice to remove extracellular fungi. THP-1 monolayers were then lysed, and fungal cells were recovered by sequential washing to a final volume of 1 mL. Recovered fungi were serially diluted and plated on YPD agar and incubated at 30⁰C for three days prior to colony-forming unit (CFU) counting. Each strain was assessed using technical replicates across three independent wells per experiment.

### Phosphoproteomics sample preparation and analysis

Wild-type and mutant *Cryptococcus neoformans* strains were grown in YPD liquid media at 30⁰C to mid-log phase. Cells were harvested by centrifugation and immediately frozen at - 80⁰C prior to analysis. Phosphoproteomic sample preparation and mass spectrometry were performed by the Duke University Proteomics and Metabolomics Core Facility using established protocols. Briefly, cell pellets were lysed by bead beating in urea-containing buffer, and total protein concentrations were determined by Bradford assay. Equal amounts of protein from each sample were reduced, alkylated, and digested with sequencing-grade trypsin using S-Trap microcolumns. Phosphopeptides were enriched by titanium dioxide (TiO_2_) affinity chromatography, followed by C18 cleanup prior to mass spectrometry. Quantitative LC-MS/MS analysis was performed on a Thermo Orbitrap Astral mass spectrometer coupled to a Vanquish Neo UPLC system using data-independent acquisition (DIA). Peptides were separated by reversed-phase chromatography and analyzed under high-resolution conditions. Raw data were processed and quantified using Spectronaut software, with peptide identification performed against the *C. neoformans* reference proteome and common contaminant databases. Phosphopeptide identifications were filtered to a 1% false discovery rate, and only phosphosites with high localization confidence were retained. Normalized phosphopeptide intensities were used to calculate fold changes between wild-type and mutant strains. Phosphopeptides were considered significantly differentially phosphorylated if they were confidently localized, detected in all three biological replicates within a group, and met thresholds of |log_2_ fold change| ≥ 1.5 with an adjusted p-value ≤ 0.05. For *mob3*Δ mutant samples, which displayed fewer phosphorylation changes relative to wild-type, phosphopeptides meeting criteria of |log_2_ fold change| ≥ 1.0 and adjusted p-value ≤ 0.1 were included in downstream analyses (Table S5).

Principal component analysis (PCA) was performed on z-score-normalized phosphopeptide expression values to assess global relationships among samples. Volcano plots were generated by collapsing phosphopeptides to unique phosphoproteins and plotting log_2_ fold change versus -log_10_ adjusted p-value for all reproducibly detected, confidently localized proteins. For Venn diagram analyses, phosphoproteins were classified as hyper- or hypophosphorylated relative to wild type based on fold change direction and significance thresholds. Because proteins may contain multiple regulated phosphosites, a phosphoprotein could be represented in both the hyper- and hypophosphorylated protein sets. Pairwise similarity between phosphoprotein sets was quantified using Jaccard similarity coefficients and visualized as a heatmap. Gene Ontology (GO) terms were assigned and enrichment analysis was performed using the built-in Gene List Analysis tools in FungiDB (https://fungidb.org), using combined sets of significantly differentially phosphorylated proteins identified in at least two STRIPAK mutant backgrounds. Enriched GO term categories were visualized as bubble plots. For protein-protein interaction network analysis, CNAG identifiers from the H99 reference genome were first converted to corresponding gene identifiers in the JEC21 reference genome, and the resulting gene lists were analyzed using STRING (https://string-db.org) with a high-confidence interaction threshold (combined score ≥ 0.70), restricting evidence to experimentally supported and curated database interactions. Network visualizations and annotations were performed using Cytoscape (version 3.10). Proteins in figures and network visualizations were labeled using the *C. neoformans* H99α nomenclature when available, or by the closest *S. cerevisiae* ortholog as determined by FungiDB or STRING.

### Statistical analysis

All statistical analyses were performed using GraphPad Prism (version 10.6). Data are presented as mean ± SEM unless otherwise indicated. Statistical significance was assessed using unpaired two-tailed Student’s *t* tests for pairwise comparisons or by one-way or two-way ANOVA followed by appropriate multiple-comparisons-corrected post hoc tests, as specified in the figure legends. For RT-qPCR, statistical analyses were performed on ΔCt values. For measurements obtained from individual cells, statistical analyses were performed using biological replicate means rather than individual cell measurements. Survival curves were analyzed using the log-rank (Mantel-Cox) test. All reported *P* values reflect correction for multiple comparisons where applicable. Statistical significance is indicated as: *P* < 0.05 (\**), P < 0.01 (**), P < 0.001 (\*\*\**), *P* < 0.0001 (****).

## Ethics statement

All animal experiments in this manuscript were approved by the Duke University Institutional Animal Care and Use Committee (IACUC) (protocol #A098-22-05). Animal care and experiments were conducted according to IACUC ethical guidelines.

## Data availability

Raw sequence reads generated from samples used in this study are available from the National Center for Biotechnology Information Sequence Read Archive (SRA) under BioProject accession number PRJNA1397472.

## Acknowledgements

We thank our laboratory manager Anna Floyd Averette for constant support, Dr. Bin Li for flow cytometry analysis, Dr. Devi Swain Lenz and the Duke Sequencing and Genomic Technologies Shared Resource for assistance with sequencing, and Dr. Erik Soderblom and the Duke Proteomics and Metabolomics Core Facility for mass spectrometry and proteome analysis. We also thank Dr. Rebecca Bacon and the Research Animal Pathology Service for histopathology assessment and assistance. PPP was supported by NIH/NIAID T32 grant AI052080-21 as a Tri-I MMPTP fellow. This work was also supported by NIH/NIAID R01 grants AI039115-28, AI050113-20, and AI172451-03 awarded to JH. JH is a co-director and fellow of the CIFAR Fungal Kingdom: Threats and Opportunities program. This research was also supported by the National Research Foundation of Korea (NRF) grant funded by the Korea government (MSIT; RS-2025-18362970, RS-2025-02215093, RS-2025-00555365) awarded to YSB.

## Author contributions

Conceptualization: Patricia P. Peterson.

Formal analysis: Patricia P. Peterson.

Funding acquisition: Yong-Sun Bahn, Joseph Heitman.

Investigation: Patricia P. Peterson, Sarah Croog, Yeseul Choi, Jin-Tae Choi.

Project administration: Yong-Sun Bahn, Joseph Heitman.

Supervision: Joseph Heitman.

Visualization: Patricia P. Peterson.

Writing – original draft: Patricia P. Peterson.

Writing – review & editing: Patricia P. Peterson, Sarah Croog, Yeseul Choi, Jin-Tae Choi, Yong-Sun Bahn, Joseph Heitman.

