## Supplemental figures for "Distinct STRIPAK subunits drive conserved and subunit-specific signaling programs in *Cryptococcus neoformans*"

**S1 Table. Strains used in this study.**

| Strain name | Description | Source/Reference |
|---|---|---|
| H99α | Wild-type *MAT*α | [1] |
| KN99**a** | Wild-type *MAT***a** | [2] |
| MCD16 | H99α *lac1*Δ*::URA5* | [3] |
| DTY1020 | H99α *cbi1*Δ*::HYG ctr4*Δ*::NAT* | [4] |
| CnLC6683 | Wild-type diploid KN99**a**/KN99α | [5] |
| YSB11754 | H99α *far9*Δ*::NAT-1* | This study |
| YSB11755 | H99α *far9*Δ*::NAT-2* | This study |
| YSB11685 | H99α *far11*Δ*::NAT-1* | This study |
| YSB11686 | H99α *far11*Δ*::NAT-2* | This study |
| YSB11687 | H99α *far11*Δ*::NAT-3* | This study |
| YSB11688 | H99α *far11*Δ*::NAT-4* | This study |
| YSB11689 | H99α *far11*Δ*::NAT-5* | This study |
| JOHE24584 | YSB11754::*NEO-GFP-FAR9-1* | This study |
| JOHE24585 | YSB11755::*NEO-GFP-FAR9-2* | This study |
| PP145 | YSB11685::*FAR11-NEO-1* | This study |
| PP113 | *MAT*α *mob3*Δ*::NAT-1* | [5] |
| PP115 | *MAT*α *mob3*Δ*::NAT-2* | [5] |
| PP117 | *MAT***a** *mob3*Δ*::NAT-3* | [5] |
| PP114 | *MAT***a** *mob3*Δ*::NAT-4* | [5] |
| PP54 | *MAT*α *pph22*Δ*::NAT* | [5] |
| PP55 | *MAT*α *pph22*Δ*::NAT* | [5] |
| PP56 | *MAT***a** *pph22*Δ*::NAT* | [5] |
| YSB9100 | *MAT*α *far8*Δ*::NAT* | [5] |
| YSB9102 | *MAT*α *far8*Δ*::NAT* | [5] |
| YSB9103 | *MAT*α *far8*Δ*::NAT* | [5] |
| JOHE24586 | YSB11754 *far9*Δ*::NAT* mouse #1 lung isolate 1 | This study |
| JOHE24587 | YSB11754 *far9*Δ*::NAT* mouse #1 lung isolate 2 | This study |
| JOHE24588 | YSB11754 *far9*Δ*::NAT* mouse #2 lung isolate 1 | This study |
| JOHE24589 | YSB11754 *far9*Δ*::NAT* mouse #2 lung isolate 2 | This study |
| JOHE24590 | YSB11755 *far9*Δ*::NAT* mouse #3 lung isolate 1 | This study |
| JOHE24591 | YSB11755 *far9*Δ*::NAT* mouse #3 lung isolate 2 | This study |
| JOHE24592 | YSB11754 *far9*Δ*::NAT* mouse #1 brain isolate 1 | This study |
| JOHE24593 | YSB11754 *far9*Δ*::NAT* mouse #1 brain isolate 2 | This study |
| JOHE24594 | YSB11754 *far9*Δ*::NAT* mouse #2 brain isolate 1 | This study |
| JOHE24595 | YSB11754 *far9*Δ*::NAT* mouse #2 brain isolate 2 | This study |
| JOHE24596 | YSB11755 *far9*Δ*::NAT* mouse #3 brain isolate 1 | This study |
| JOHE24597 | YSB11755 *far9*Δ*::NAT* mouse #3 brain isolate 2 | This study |

**S2 Table. Primers used in this study.**

| Primer # | Sequence | Purpose |
|---|---|---|
| M13F | GTAAAACGACGGCCAG | To amplify drug resistance cassettes |
| M13R | CAGGAAACAGCTATGAC | To amplify drug resistance cassettes |
| JOHE52463 | CTGGCGGAGGATAGAAGC | *ACT1* promoter reverse primer |
| JOHE52464 | GCGAATTCGAGACAGACATCG | *TRP1* terminator forward primer |
| B21774 | TTATCGTCCGTCTTCGATCC | *FAR9* deletion |
| B21775 | TCACTGGCCGTCGTTTTACGCTTGTCCTTGTTGGGAGAG | |
| B21776 | CATGGTCATAGCTGTTTCCTGTGGTCGTGGCGATTGTAGTA | |
| B21777 | CTCCCTACGCTTCATCTCCA | |
| B21510 | CTGATGGGGATTTGGTGAAC | *FAR11* deletion |
| B21511 | TCACTGGCCGTCGTTTTACCGGAGCGTTCTCTTGTTGAT | |
| B21512<br>B21513 | CATGGTCATAGCTGTTTCCTGGGTGCCGTGGATGAAGTAGA<br>GGTGTGGATAGGGATGAGGA | |
| JOHE52395 | TGACTCAGATGAACGCAACC | *FAR9* screening primers |
| JOHE52396 | GGGCCATTAACCATTTAGCA | |
| JOHE52429 | ACTCCTAGCAGAACCAGGAC | |
| JOHE52430 | CCTCTTCCTCCTCATATGCT | |
| JOHE52381 | CGTCTTCTCTCGCTTCCATC | *FAR11* screening primers |
| JOHE52382 | GTACCAGGAGATCCCGAACA | |
| B21514 | CTGGAGTTCTCGGTCCTCTG | |
| B21843 | CCTGCTCCCAAACGTATCAT | |
| JOHE52417 | GTGAAAGATGGCAAAACTCACA | *FAR11::NEO* complementation |
| JOHE56876 | TTGCGGCCGCAGGTGGCGGTGG | |
| JOHE56886 | CACACTGGCGGCCGTTACTAATTTTTAATCGATGCGCTCA | |
| JOHE56887 | CCACCGCCACCTGCGGCCGCCTCTCCGTAAGTATATTCTA | |
| JOHE56885 | ACCGATCATACCCGCCAAAG | |
| JOHE55518 | TAGTAACGGCCGCCAGTGTGCTGG | |
| JOHE56283 | AACGGCCGCCAGTGTGCTGGAATTCGCTGCGAGGATGTGAGCTG | *FAR9::NEO* complementation |
| JOHE56284 | AGTGTGATGGATATCTGCAGAATTCGGTTTATCTGTATTAACACGGAAGAGATG | |
| JOHE56351 | CGTGCCGTTTGTGTAAGAGA | |
| JOHE56352 | GTCATAGCTGTTTCCTGTACTTCCTTTGATGTGTCCTCG | |
| JOHE56355 | CACTGGCCGTCGTTTTACAGTCACATGTAACTTAATC | |
| JOHE56356 | GGTGCTATCATGGCTTTGGT | |
| JOHE56353 | GACACATCAAAGGAAGTACAGGAAACAGCTATGAC | |
| JOHE56354 | GATTAAGTTACATGTGACTGTAAAACGACGGCCAGTG | |
| JOHE57898 | ATGCTCCTATGTTTGTCTGC | *GAPDH* qPCR |
| JOHE57899 | GCAGTAGTAGCATGGACAG | |
| JOHE57900 | ATGATGCAAGGAGTGTGGCA | *FAR9* qPCR |

| | | |
|---|---|---|
| JOHE57901 | GCTTCAAGGTCCTTAGGCGT | |
| JOHE57902 | CATTCCCCCTGCTCCACTTT | *FAR11* qPCR |
| JOHE57903 | TGGTGCCGCTAGTAGAGTCT | |

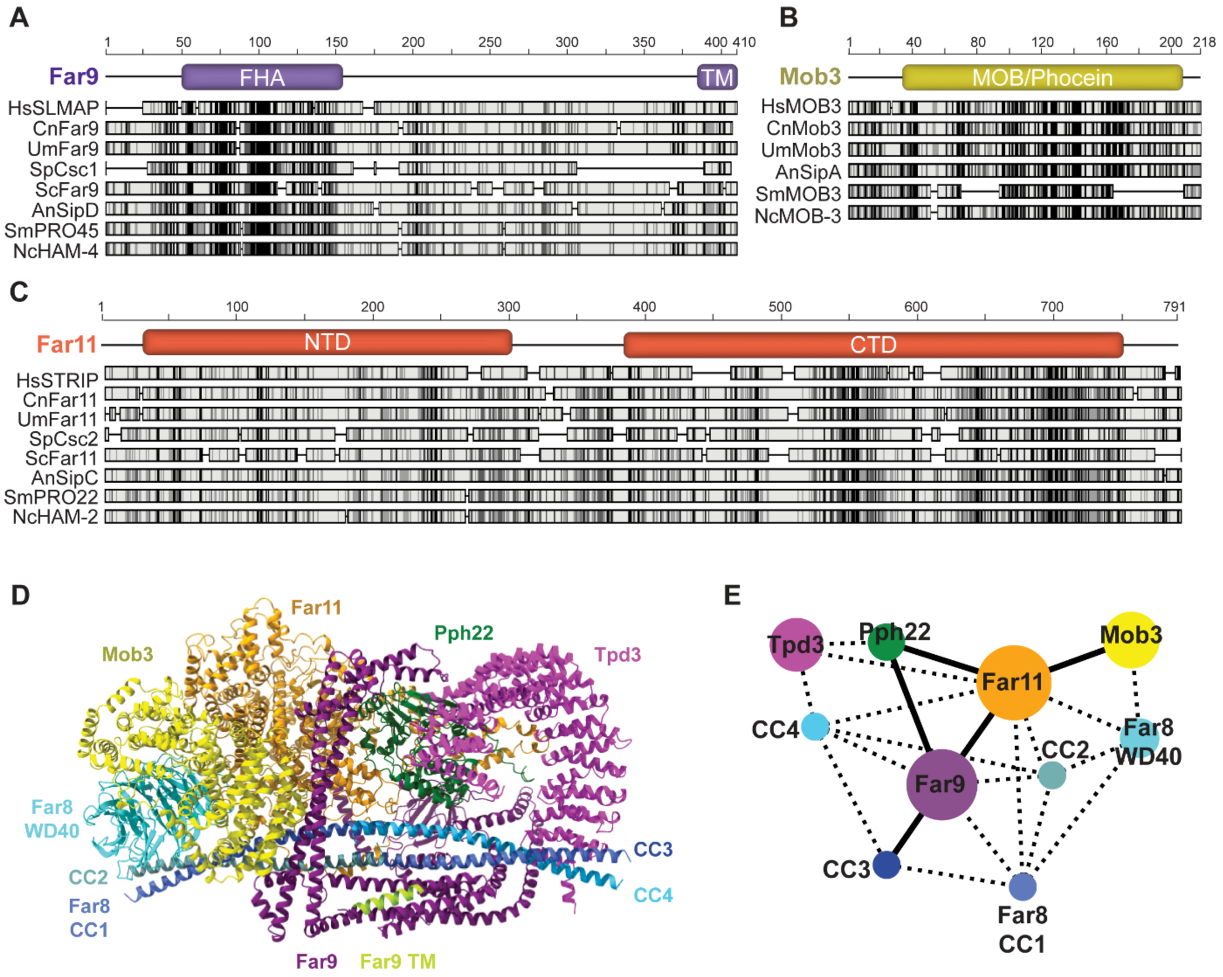


**S1 Fig. Conserved domain architecture but divergent primary sequence among STRIPAK subunits across eukaryotes.**

Multiple sequence alignments of STRIPAK components Far9 (A), Mob3 (B), and Far11 (C) from representative fungi and humans reveal distinct conservation patterns. Far9 retains a conserved FHA domain and C-terminal transmembrane (TM) region, Mob3 exhibits strong conservation throughout the MOB/phocein domain, and Far11 displays conserved N-terminal and C-terminal regions, despite substantial overall sequence divergence. Alignments were generated using MAFFT and colored based on amino acid similarity using the BLOSUM62 matrix, with darker shading indicating higher conservation. Species abbreviations: Hs, *Homo sapiens;* Cn, *Cryptococcus neoformans*; Um, *Ustilago maydis*; Sp, *Schizosaccharomyces pombe*; Sc, *Saccharomyces cerevisiae*; An, *Aspergillus nidulans*; Sm, *Sordaria macrospora*; Nc, *Neurospora crassa.* (D) AlphaFold3 multimer prediction of the six-subunit *Cn*STRIPAK complex (ipTM= 0.51, pTM= 0.56) [6]. The predicted stoichiometry of the Far8 subunit is shown [7], with one WD40 β-propeller and four coiled-coil (CC) domains included in the model. The TM domain of the tail-anchored subunit Far9 is also highlighted (GRAVY= 2.59). (E) Predicted interaction interfaces among STRIPAK subunits, highlighting Far11 serves as the central organizing scaffold with eight predicted interaction surfaces. Structural analyses and visualizations were performed in ChimeraX [8].

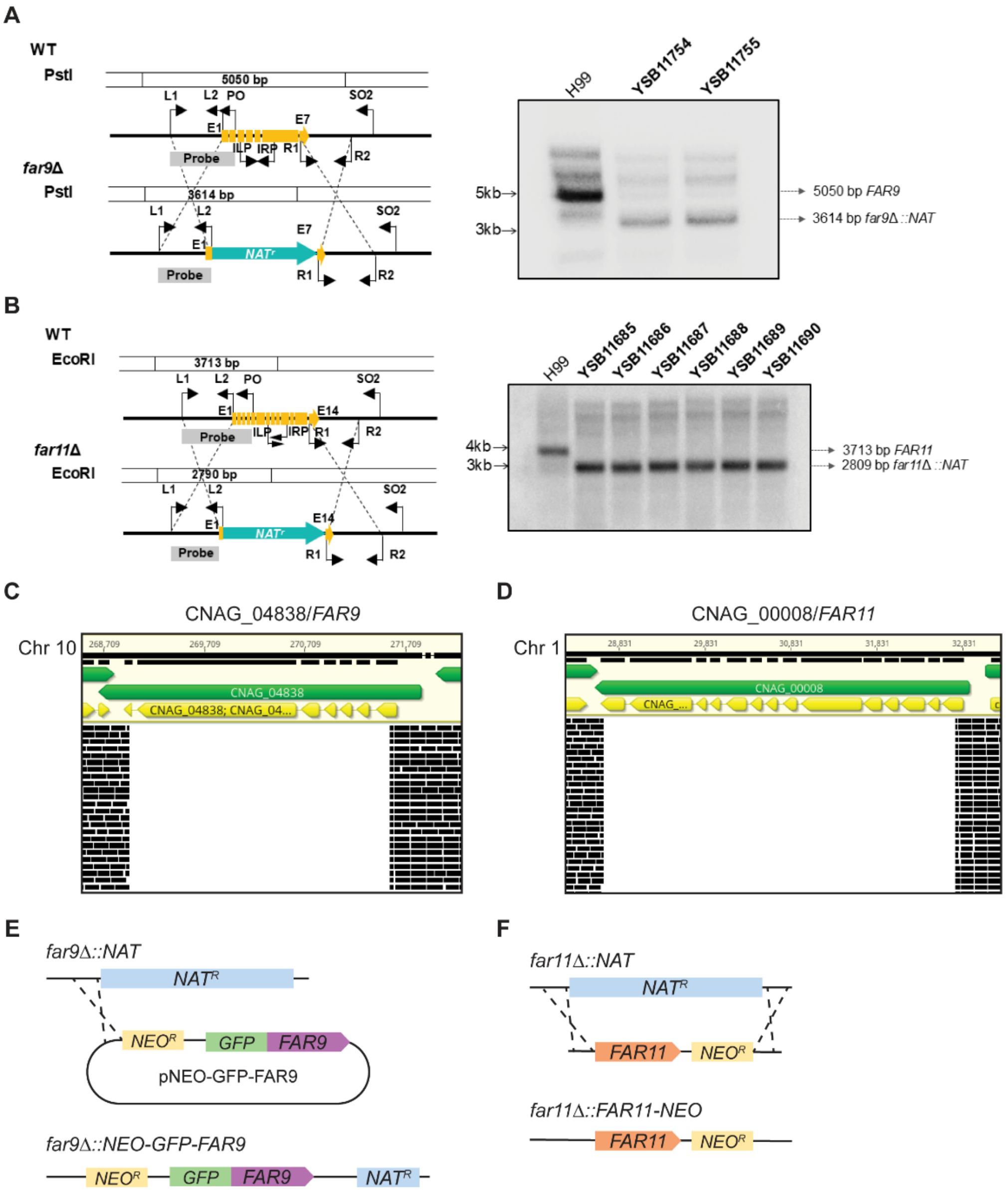


**S2 Fig. *far9*Δ and *far11*Δ strain construction and confirmation of deletion mutants.** Mutant strain construction design for (A) *far9*Δ and (B) *far11*Δ in the H99α background, along with Southern blot analysis confirming deletion of the target gene. Representative snapshots of Illumina whole genome sequencing analysis in (C) *far9*Δ (YSB11754) and (D) *far11*Δ (YSB11685) mutant strains showing the absence of reads mapping to the *FAR9* or *FAR11* open reading frames, respectively. The coding sequences (CDS) of each gene are depicted in yellow, at the indicated chromosome coordinates, and individual DNA reads mapping to the H99α reference sequence are shown beneath in black. (E) Schematic of the *far9*Δ::*NEO-GFP-FAR9* complementation strategy. A GFP-tagged *FAR9* construct carrying the *NEO* selectable marker was integrated by homologous recombination upstream of the *far9*Δ*::NAT* deletion cassette, preserving the native *FAR9* promoter and 3′ UTR. (F) Schematic of the *far11*Δ*::FAR11-NEO* complementation strategy. The *FAR11* coding sequence together with the *NEO* selectable marker was reintroduced by homologous recombination at the endogenous *FAR11* locus, replacing the *far11*Δ*::NAT* deletion cassette while preserving the native promoter and 3′ UTR.

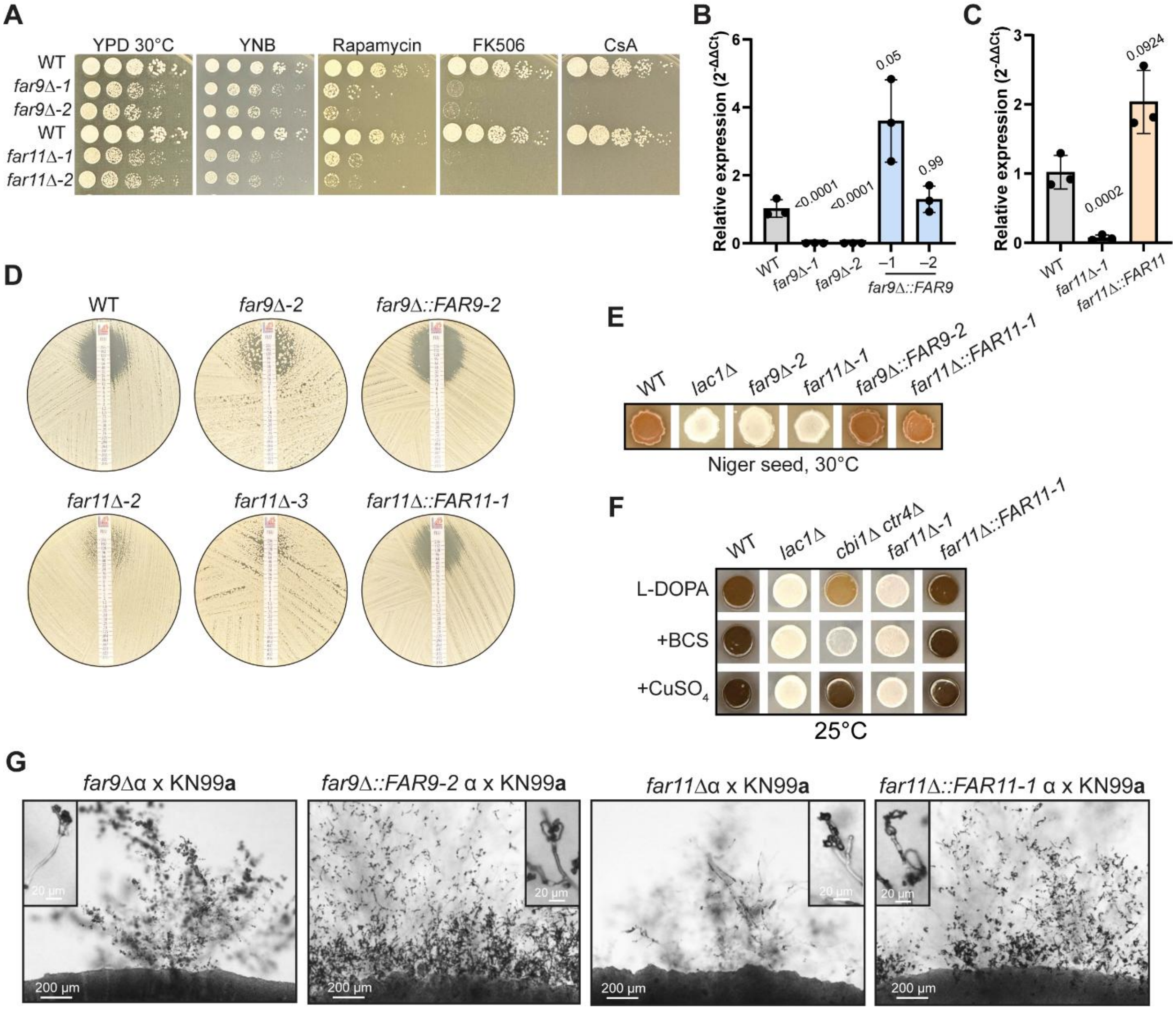


**S3 Fig. *far9*Δ and *far11*Δ growth phenotypes under nutrient and drug stress, and during sexual development.**

(A) Wild-type (H99α), *far9*Δ, and *far11*Δ were serially diluted and spotted onto YPD, YNB, and YPD supplemented with rapamycin (100 ng/mL), FK506 (1 µg/mL), or cyclosporine A (CsA; 100 µg/mL). Plates were incubated at 30°C and images after three days. (B and C) Relative *FAR9* (B) and *FAR11* (C) transcript abundance measured by RT-qPCR in WT, deletion mutants, and complemented strains. Gene expression was normalized to *GAPDH* and is presented as $2^{\wedge-\Delta\Delta Ct}$ relative to WT. Bars represent the mean ± SD of three biological replicates, with each point representing the mean of two technical replicates from an individual biological replicate. Statistical analyses were performed on normalized ΔCt values using one-way ANOVA with Dunnett's multiple-comparisons test. (D) Fluconazole Etest drug susceptibility assay of independent *far9*Δ and *far11*Δ mutant isolates along with the *FAR9* and *FAR11* complemented strains. Representative plates from the same assay shown in Figure 1B are displayed. (E) Melanin production on Niger seed medium at 30°C by WT, deletion mutants, and complemented strains. Representative images are shown from the assay described in Figure 1C. (F) Melanin production on L-DOPA medium at 25°C under standard conditions or following supplementation

with the copper chelator bathocuproine disulfonate (BCS) or $CuSO_4$. Representative images demonstrate restoration of the *far11*Δ melanization phenotype following complementation.
(G) Representative images of sexual crosses between WT and independent clones of *far9*Δ and *far11*Δ mutants, as well as the corresponding complemented strains, demonstrating that reintroduction of the WT allele restores normal sexual development. Images are from the same experiment shown in Figure 1D.

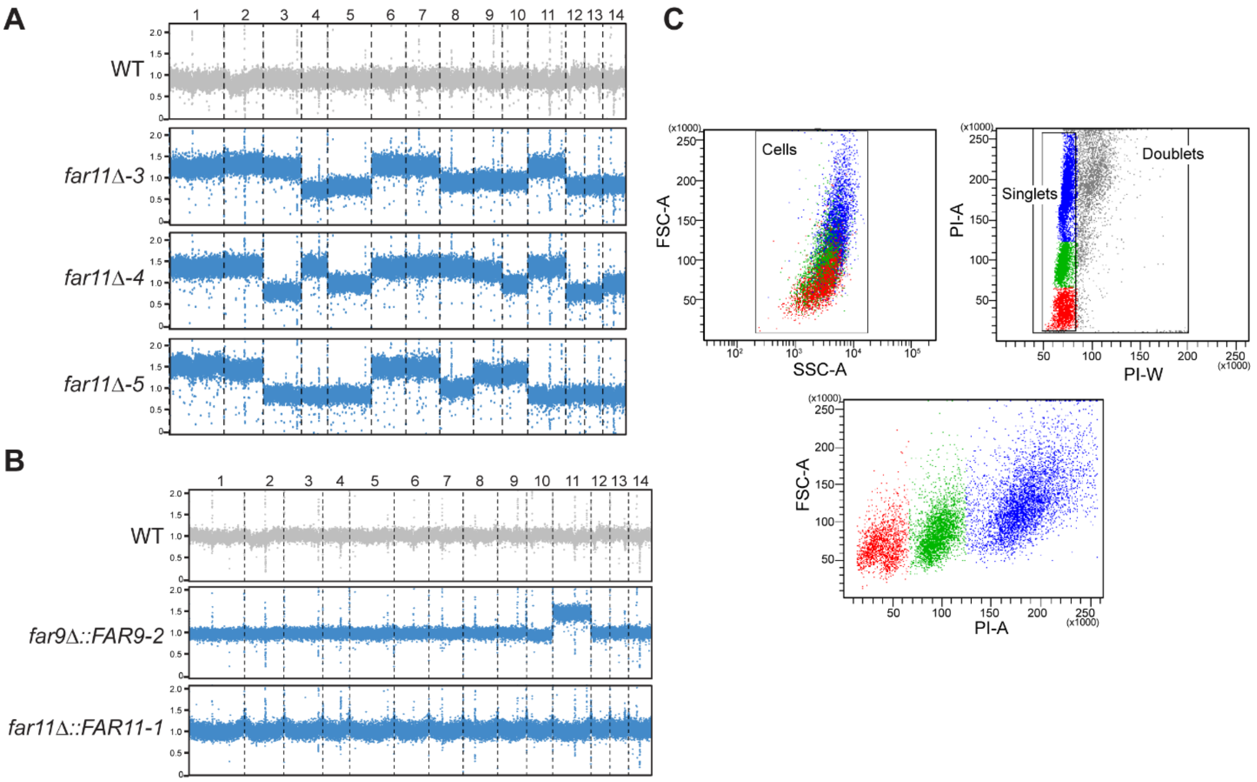


D

| | 1C P1 | 2C P2 | 4C P3 | >4C P4 |
|---|---|---|---|---|
| WT | 55.5 | 40.8 | 3.5 | 0.2 |
| WT/WT | 3.9 | 64.3 | 29.5 | 2.3 |
| *far9Δ-1* | 13.7 | 40.6 | 32.3 | 13.4 |
| *far9Δ-2* | 5.2 | 40.1 | 37.5 | 17.2 |
| *far11Δ-1* | 2.9 | 35.8 | 43.2 | 18.1 |
| *far11Δ-2* | 4.8 | 23.7 | 48.9 | 22.6 |
| *far11Δ-3* | 5.8 | 31.7 | 45.4 | 17.1 |

YPD, 30°C

| | 1C P1 | 2C P2 | 4C P3 | >4C P4 |
|---|---|---|---|---|
| WT | 41.5 | 32.6 | 21.9 | 4.0 |
| WT/WT | 9.2 | 44.9 | 30.4 | 15.5 |
| *far9Δ-1* | 6.9 | 35.5 | 30.0 | 27.6 |
| *far9Δ-2* | 7.2 | 43.2 | 25.5 | 24.1 |
| *far11Δ-1* | 6.3 | 14.0 | 55.4 | 24.3 |
| *far11Δ-2* | 8.1 | 6.2 | 42.9 | 42.8 |
| *far11Δ-3* | 6.3 | 6.2 | 57.1 | 30.4 |

10% serum, 37°C

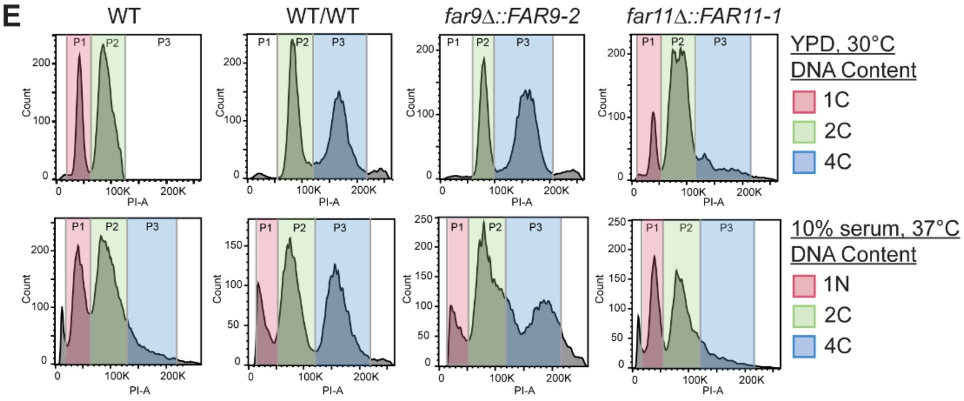

**S4 Fig. Quantification of DNA content distribution in *far9*Δ and *far11*Δ mutants.**
(A) Read-depth analysis from whole-genome sequencing of additional independent *far11*Δ mutant isolates. Read coverage was normalized to the genome-wide median. (B) Read-depth analysis from whole-genome sequencing of *far9*Δ*::FAR9* and *far11*Δ*::FAR11* complemented strains. Read coverage was normalized to the genome-wide median. Complementation of *FAR11* restored a euploid genome profile, whereas the *far9*Δ*::FAR9* strain retained increased chromosome 11 copy number. (C) Flow-cytometry gating strategy used for DNA-content analysis. Debris was excluded based on forward- and side-scatter properties (FSC-A vs. SSC-A). Single cells were then identified by pulse-processing parameters (PI-A vs. PI-W), and events with increased pulse width consistent with doublets or cell aggregates were excluded. DNA-content populations were subsequently defined using propidium iodide fluorescence intensity (PI-A). High-DNA-content events retained singlet pulse characteristics while exhibiting increased PI-A and FSC-A, consistent with enlarged single cells rather than multicellular aggregates. (D) Percentages of cells within each DNA content gate (P1–P4) corresponding to 1C, 2C, 4CN, and >4C populations were calculated from flow cytometry analyses shown in Figure 2B and 2C. Strains were grown in YPD at 30°C or 10% serum at 37°C, as indicated. Values represent the proportion (%) of total events falling within each gate for the indicated strain. (E) Flow-cytometry analysis of DNA content in WT (H99α), WT/WT (KN99α/**a**), *far9*Δ*::FAR9*, and *far11*Δ*::FAR11* complemented strains grown in YPD at 30°C or 10% serum at 37°C. Cells were stained with propidium iodide, and peaks corresponding to 1C, 2C, and 4C DNA content are indicated.

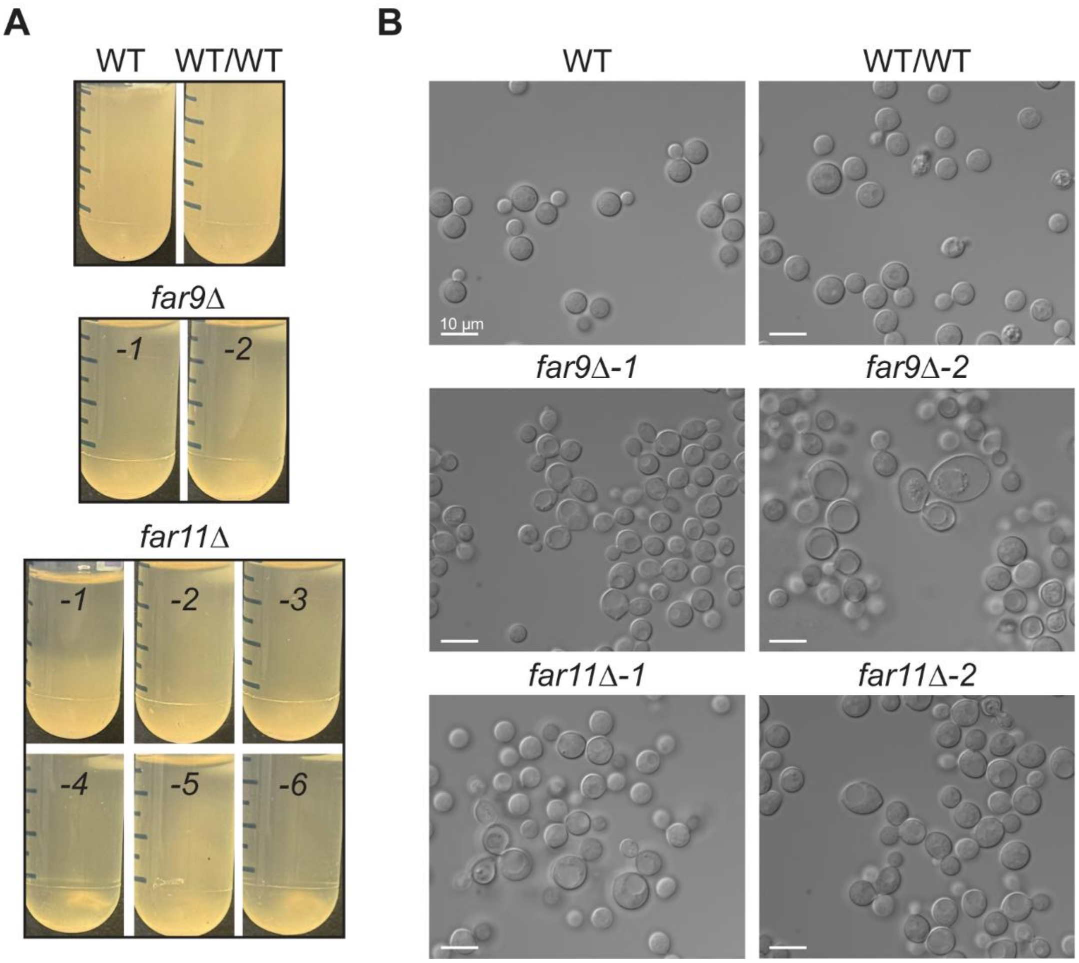


**S5 Fig. Altered settling behavior and aggregation of *far9*Δ and *far11*Δ mutant cells in liquid medium.**

(A) Representative images of liquid YPD cultures of wild-type haploid (WT), diploid (WT/WT), *far9*Δ, and *far11*Δ strains grown overnight at 30°C. Cultures were normalized to the same optical density, vortexed to resuspend cells, and then allowed to remain undisturbed at room temperature for 1 hour prior to imaging. Under these conditions, *far9*Δ and *far11*Δ mutants exhibited increased sedimentation and visible cell clustering relative to wild-type controls. (B) Differential interference contrast (DIC) microscopy images of cells taken from the bottom 1 mL of cell culture depicted in (A) show increased aggregation and heterogeneity in cell size in *far9*Δ and *far11*Δ mutants compared to WT controls. Scale bars represent 10 μm.

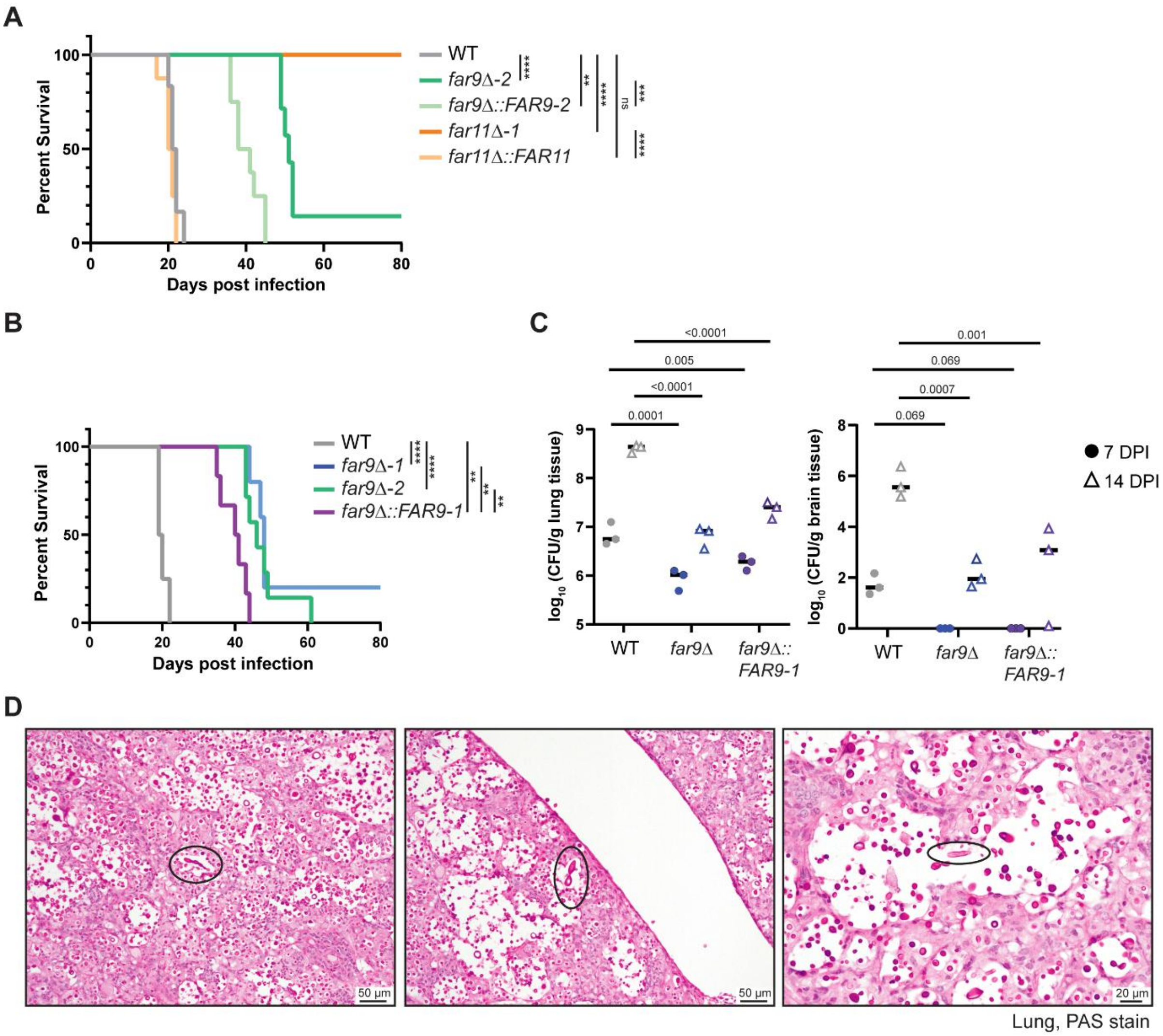


**S6 Fig. Survival and fungal burden in mice infected with WT, *far9*Δ, *far9*Δ*::FAR9* strains.** (A) Survival analysis of A/J mice infected with WT(H99α), *far9*Δ, *far11*Δ, and the corresponding complemented strains, demonstrating restoration of WT virulence in the *far11*Δ*::FAR11* strain and partial restoration in the *far9*Δ*::FAR9* strain (1 x $10^5$ cells/mouse; n=10 per group). Statistical significance was determined using the log-rank (Mantel-Cox) test, with comparisons shown relative to the WT H99α strain. (B) Survival analysis of A/J mice infected with WT, *far9*Δ or the second independently generated *far9*Δ*::FAR9* complemented strain. This experiment was performed independently from the survival and fungal burden analyses shown in Figure 4A and 4B. (C) Fungal burden, expressed as CFUs per gram of tissue, recovered from lungs and brains at 7 and 14 days post-infection (DPI). Both *far9*Δ and *far9*Δ*::FAR9* strains exhibited significantly reduced lung fungal burdens at 7 and 14 DPI compared with WT. In the brain, neither *far9*Δ nor *far9*Δ*::FAR9* was detectable at 7 DPI but both were recovered at 14 DPI at several orders of magnitude lower than WT, consistent with delayed dissemination. Statistical significance was assessed using two-way ANOVA with Dunnett's multiple-comparisons test. (D) Other atypical cell morphologies (black circles), including pseudohyphae and elongated cells, are observed in infections with *far9*Δ mutants in the lungs. Tissues were collected for analysis at the termination of infection.

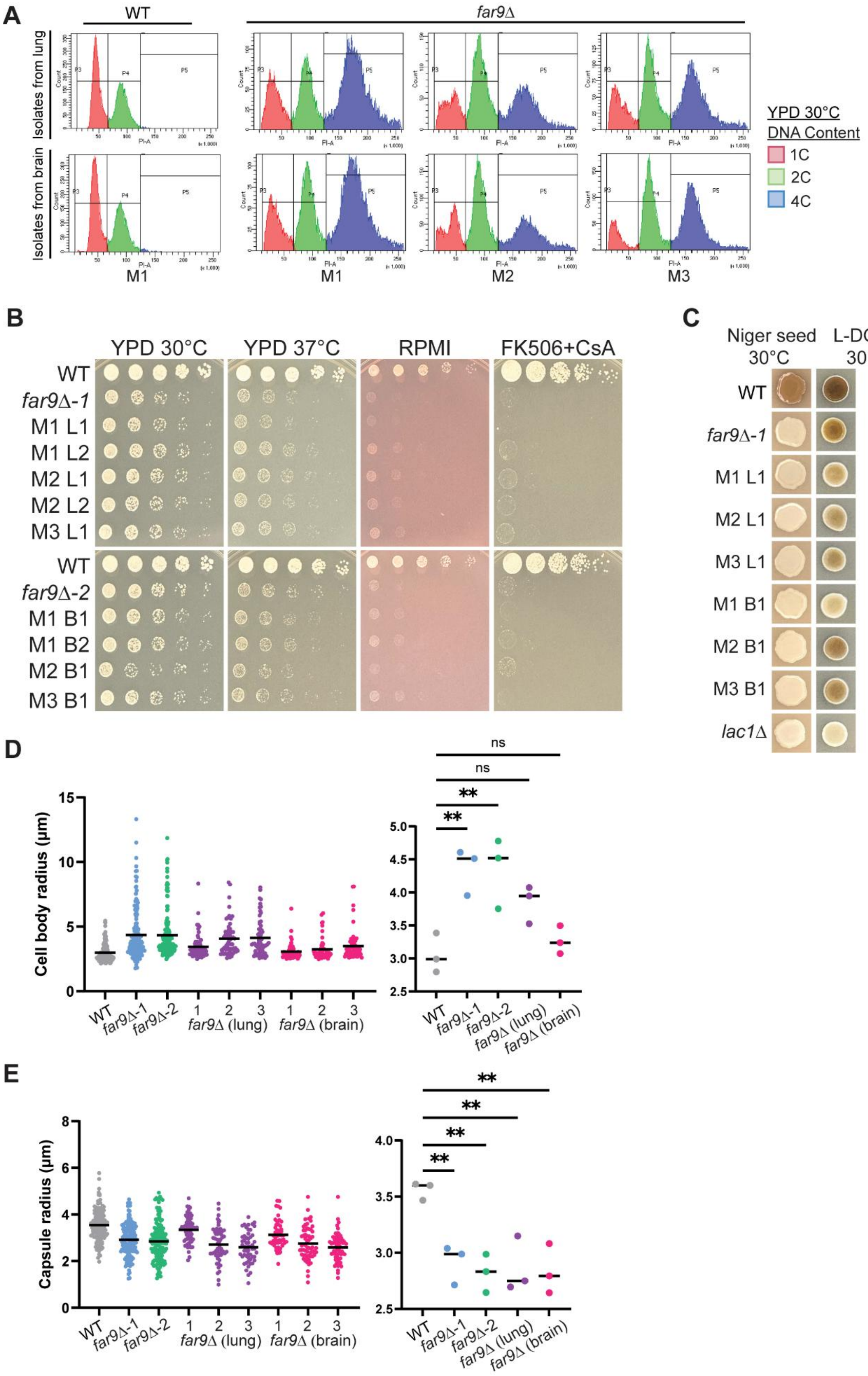
A
WT
far9Δ
Isolates from lung
Isolates from brain
M1
M1
M2
M3
YPD 30°C
DNA Content
1C
2C
4C
B
YPD 30°C
YPD 37°C
RPMI
FK506+CsA
WT
far9Δ-1
M1 L1
M1 L2
M2 L1
M2 L2
M3 L1
WT
far9Δ-2
M1 B1
M1 B2
M2 B1
M3 B1
C
Niger seed 30°C
L-DOPA 30°C
WT
far9Δ-1
M1 L1
M2 L1
M3 L1
M1 B1
M2 B1
M3 B1
lac1Δ
D
Cell body radius (μm)
WT
far9Δ-1
far9Δ-2
far9Δ (lung)
far9Δ (brain)
ns
ns
**
**
E
Capsule radius (μm)
WT
far9Δ-1
far9Δ-2
far9Δ (lung)
far9Δ (brain)
**
**
**
**

**S7 Fig. Phenotypic characterization of *far9*Δ isolates recovered from murine infection.** (A) Flow-cytometry analysis of DNA content in WT and far9Δ isolates recovered from the lungs and brains of infected animals. Cells were grown in YPD at 30°C, stained with propidium iodide, and analyzed by flow cytometry. Peaks corresponding to 1C, 2C, and 4C DNA content are indicated. M1-M3 denote independent mice. (B) WT (H99α), *far9*Δ parental strains, and *far9*Δ mutant isolates from the brain and lungs of independent animals were serially diluted ono YPD at 30ºC, YPD at 37ºC, RPMI at 30ºC, and YPD supplemented with FK506 (1 μg/mL) and CsA (100 μg/mL) at 30ºC. Plates were imaged after two days of incubation. Shown are the same isolates described in Figure 4. M1-M3 denote independent mice; L, lung isolate; B, brain isolate. (C) Melaninization of strains on Niger seed or L-DOPA medium. Plates were incubated for two days prior to imaging. Mouse-passaged *far9*Δ isolates, like the *far9*Δ parental strain, failed to produce melanin on Niger seed media. On L-DOPA recovered isolates displayed melanin production comparable to, or modestly reduced, relative to the parental strain. H99α and a *lac1*Δ mutant were included as positive and negative controls, respectively. (D and E) Quantification of cell-body radius (D) and capsule radius (E) of WT, parental *far9*Δ strains, and mouse-passaged *far9*Δ isolates grown in 10% fetal bovine serum at 37°C with 5% $CO_2$ and stained with India ink. Scatter plots (left) show individual cells, whereas plots on the right show the mean values from three biological replicates (WT and parental *far9*Δ strains) or independent recovered isolates. Statistical significance was determined using one-way ANOVA with Dunnett's multiple-comparisons test.

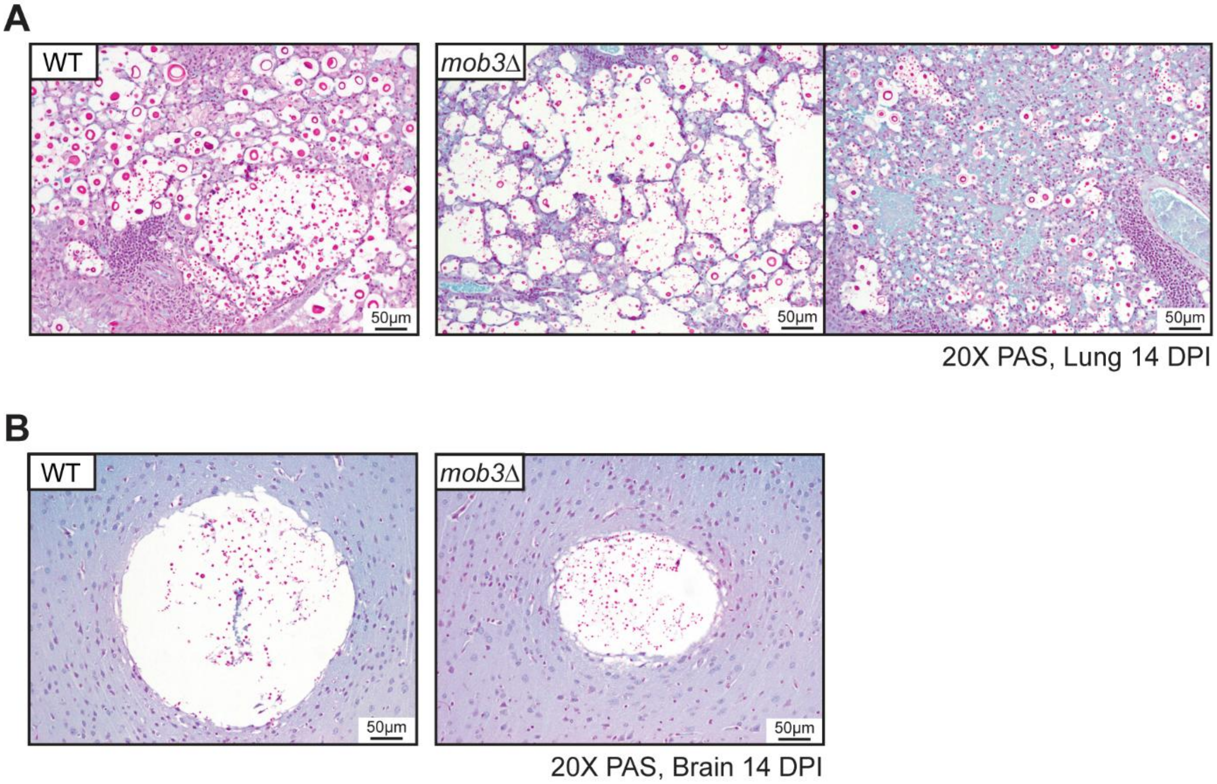


**S8 Fig. Histopathological analysis of lung and brain tissues from WT- and *mob3*Δ-infected mice.**

(A) Representative PAS-stained lung sections from WT and the independent *mob3*Δ-1 and *mob3*Δ-2 isolates at 14 days post-infection. Together with the *mob3*Δ-3 images shown in Figure 6F, all three independent *mob3*Δ isolates exhibited overall smaller fungal cells *in vivo*. (B) Representative PAS-stained brain sections from WT and the *mob3*Δ-3 isolate at 14 days post-infection. All three independent *mob3*Δ isolates exhibited similar brain pathology, characterized by relatively homogeneous fungal populations, comparable cryptococcoma formation, and minimal surrounding host inflammation. PAS, Periodic Acid-Schiff.
